# Codon Pairs are Phylogenetically Conserved: Codon pairing as a new class of phylogenetic characters

**DOI:** 10.1101/654947

**Authors:** Justin B. Miller, Lauren M. McKinnon, Michael F. Whiting, Perry G. Ridge

## Abstract

Identical codon pairing and co-tRNA codon pairing increase translational efficiency within genes when two codons that encode the same amino acid are located within a ribosomal window. By examining both identical and co-tRNA codon pairing across 23 423 species, we determined that both pairing techniques are phylogenetically informative across all domains of life using either an alignment-free or parsimony framework. We also determined that conserved codon pairing typically has a smaller window size than the length of a ribosome. We also analyzed frequencies of codon pairing for each codon to determine which codons are most likely to pair. The alignment-free method does not require orthologous gene annotations and recovers species relationships that are comparable to other alignment-free techniques. Parsimony generally recovers phylogenies that are more congruent with the established phylogenies than the alignment-free method. However, four of the ten taxonomic groups do not have sufficient ortholog annotations and are therefore recoverable using only the alignment-free methods. Since the recovered phylogenies using only codon pairing largely match established phylogenies and are comparable to other algorithms, we propose that codon pairing biases are phylogenetically conserved and should be considered in conjunction with current techniques in future phylogenomic studies. Furthermore, the phylogenetic conservation of codon pairing indicates that codon pairing plays a greater role in the speciation process than previously acknowledged.

**Availability:** All scripts used to recover and compare phylogenies, including documentation and test files, are freely available on GitHub at https://github.com/ridgelab/codon_pairing.

## Introduction

Phylogenies allow biologists to infer similar characteristics of closely related species and provide an evolutionary framework for analyzing biological patterns (Soltis and Soltis 2003). Phylogenies are statements of homology, and represent a continuity of biological information (Haszprunar 1992). Although genetic data allow researchers to analyze more species cheaper and faster than morphological features, molecular data typically require data cleaning (e.g., alignment, annotation, and ortholog identification) before they become useful (Philippe, et al. 2011). After orthologs are identified, phylogenies can be recovered through parsimony (Farris 1983; Wilgenbusch and Swofford 2003), maximum likelihood (Felsenstein 1981), Bayesian inference (Yang and Rannala 2012), or distance-based techniques such as neighbor-joining (Saitou and Nei 1987).

Although many novel phylogenetic algorithms have been developed, parsimony remains important in phylogenetic analyses because of its explanatory power, which limits the number of separate character state origins (Farris 2008). Therefore, the most parsimonious tree depicts character homology and limits *ad hoc* hypotheses. Although sequence alignments are typically used in parsimony analyses, aversion to certain codons was recently shown to be phylogenetically conserved within orthologs (Miller, et al. 2017; Miller, et al. 2019b). In those analyses, each ortholog was encoded with 64 characters, one for each codon. Codons were given a binary representation of ‘1’ if the codon was used within the ortholog and ‘0’ if the codon was not used, and each character was added to a matrix. The parsimony analysis proceeded with the encoded character matrix, recovering trees that largely resembled previously-reported phylogenies.

Alignment-free methods have recently gained traction because they do not require a sequence alignment, which allows species with unannotated genes to be placed on the species tree. Furthermore, the alignment-free algorithms are usually computationally inexpensive, which allows for more species to be compared. Proponents of alignment-free techniques claim that they are resistant to shuffling and recombination events and are not affected by assumptions regarding a high correlation between sequence changes and evolutionary time (Bonham-Carter, et al. 2014) or limited by potential errors in orthology (Zielezinski, et al. 2017). Alignment-free techniques typically use Chaos Theory to calculate distances of basic genomic features (e.g., GC content, oligomer frequency, etc.) that are then used to recover the phylogeny (Vinga and Almeida 2003; Chan, et al. 2014). More recently, another technique limits the alignment-free search space to all genic regions within a species, comparing species-wide patterns of codon aversion and amino acid aversion independent of gene annotations (Miller, et al. 2019a). Generally, alignment-free approaches can be grouped into three main types. The first group determines the frequency of words of a certain length (e.g., FFP (Sims, et al. 2009; Jun, et al. 2010) and CVTree (Zuo and Hao 2015)). The second group finds match lengths between sequences (e.g., ACS (Ulitsky, et al. 2006), KMACS (Leimeister and Morgenstern 2014), and Kr (Haubold, et al. 2009)). The last group calculates informational content between sequences (e.g., Co-phylog (Yi and Jin 2013), FSWM (Leimeister, et al. 2017), andi (Haubold, et al. 2015), and CAM (Miller, et al. 2019a)).

Here, we present two novel approaches that capitalize on biases in codon usage to determine species relationships using either a parsimony or alignment-free framework. Codons are sequences of three consecutive nucleotides of coding DNA that are transcribed into mRNA, mRNA is translated into amino acids, and amino acids form proteins (Crick 1970). The 20 canonical amino acids are formed from 61 codons, with the other three codons encoding the stop signal (Crick, et al. 1961). Although multiple codons encode the same amino acid, an unequal distribution of synonymous codons occurs within species, suggesting that synonymous codons might play different roles in species fitness (Sharp and Li 1986). An unequal distribution of tRNA anticodons directly coupling codons led to the wobble hypothesis: tRNA anticodons do not need to latch onto all three codon nucleotides for translation (Crick 1966). Codon usage is also highly associated with the most abundant tRNA present in the cell (Post, et al. 1979) and codon usage patterns affect gene expression (Gutman and Hatfield 1989).

Recharging a tRNA while the tRNA is still attached to the ribosome is used to increase translational efficiency and decrease overall resource utilization. This process occurs when codons encoding the same amino acid are located in close proximity to each other on the mRNA strand (Cannarozzi, et al. 2010). Co-tRNA codon pairing is when two non-identical codons that encode the same amino acid are near each other in a gene and the tRNA is recharged to translate both codons before the tRNA diffuses. Similarly, identical codon pairing occurs when identical codons are near each other in a gene and the tRNA is recharged to translate both codons before the tRNA diffuses. Co-tRNA and identical codon pairing conserve resources and increase translational speed by approximately 30% (Cannarozzi, et al. 2010). Co-tRNA codon pairing has previously been reported as more prominent in eukaryotes, while identical codon pairing has been reported in eukaryotes, bacteria (Shao, et al. 2012), and archaea (Zhang, et al. 2013).

Our analyses suggest that both identical codon pairing and co-tRNA codon pairing are phylogenetically conserved and prominent in all domains of life. We further show that combining the two techniques generally recovers more congruent phylogenies compared to the Open Tree of Life (OTL) (Hinchliff, et al. 2015) and the NCBI Taxonomy Browser (Sayers, et al. 2009; Sayers, et al. 2010; Sayers, et al. 2011; Sayers, et al. 2012), and comparable trees to other phylogenomic methods. Therefore, we propose that codon pairing be considered as another method to recover phylogenies.

## Results

We aimed to determine how phylogenies recovered using identical codon pairing and/or co-tRNA pairing compare to traditional methods (i.e., parsimony and maximum likelihood) and other alignment-free methods. First, we determined the theoretical maximum number of character states for each gene using codon pairing in order to determine the maximum number of species we can differentiate using this technique. For identical codon pairing, there are 61 possible pairing combinations (64 codons – 3 stop codons), meaning each gene can separate a maximum of 2^61^ = 2.306 × 10^18^ species. For co-tRNA codon pairing, there are 18 amino acids that use more than one codon, meaning there are 18 possible pairing combinations. Using co-tRNA codon paring, each gene can separate a maximum of 2^18^ = 262,144 species. Using the combined approach, there are 20 possible pairing combinations, one for each of the 20 amino acids. This approach allows each gene to separate a maximum of 2^20^ = 1,048,576 species. Since orthologous genes are conserved between species, closely related species share a higher number of codon pairings than more distantly related species. We observed this overlap in codon pairings, with closely related species often having smaller observed distances than distantly related species.

Table 1 shows the number of species that were included in each analysis after the preprocessing filters were applied (e.g., each species in the parsimony analysis included at least 5% of the parsimony-informative characters). In total, we included 23 428 species, with each species generally containing thousands of genes. Supplementary Tables 1-3 show the number of species that were included for each ribosomal window size in each of the three parsimony analyses. The alignment-free methods included all species because the method is not affected by missing orthologous gene annotations. The NCBI taxonomy contains more taxonomic relationships than the OTL, and the OTL does not contain any viruses. Furthermore, depending on the taxonomic group, the species trees vary between the OTL and the NCBI taxonomy by 1-9%, with the mammal phylogenies being the most similar and the fungi phylogenies being the least similar. Parsimony and maximum likelihood used similar numbers of species in each analysis. A stricter filter was applied to the parsimony analysis than the maximum likelihood analysis, which required the parsimony character matrix to include at least 5% of the parsimony-informative characters. After this filter was applied, we required that at least 5% of the total number of species be included in the analysis (e.g., if 100 species were analyzed, at least five species must pass the preprocessing step for the taxonomic group to be included). Applying this filter removed the results from all species, bacteria, and viruses from both the parsimony and maximum likelihood analyses. This filter also removed fungi from the parsimony analysis.

**Table 1:** Number of species passing preprocessing filters and analyzed by each algorithm.

| Taxonomic Group | Alignment-free | Parsimony | Maximum Likelihood | NCBI Taxonomy | OTL |
| --- | --- | --- | --- | --- | --- |
| All | 23 428 | 0 | 0 | 22 794 | 12 337 |
| Archaea | 418 | 100 | 418 | 416 | 362 |
| Bacteria* | 15 068 | 0 | 0 | 14 612 | 11 227 |
| Fungi | 234 | 0 | 58 | 234 | 214 |
| Invertebrates | 149 | 57 | 57 | 149 | 147 |
| Plants | 89 | 61 | 60 | 89 | 87 |
| Protozoa | 75 | 15 | 24 | 75 | 75 |
| Mammals | 107 | 97 | 100 | 107 | 105 |
| Other vertebrates | 123 | 114 | 118 | 123 | 120 |
| Viruses* | 7 233 | 0 | 0 | 7 045 | 0 |

The alignment-free methods include codon pairing, CAM, and FFP, and did not require any preprocessing. Parsimony used a stricter preprocessing cutoff than maximum likelihood, and therefore used fewer species. The NCBI taxonomy includes viruses and more species than the OTL. Zero species passed the filters when fewer than 5% of the total species had sufficient ortholog annotations to run the analysis.

After filtering for parsimony-informative codons, we used parsimony to recover phylogenies with the highest percent overlap based on codon pairings. The identical codon pairing parsimony analysis was based on 794 (invertebrates) to 197 074 (mammals) parsimony-informative codons. The co-tRNA codon pairing analysis used 382 (invertebrates) to 94 018 (mammals) parsimony-informative codons. The combined codon pairing analysis used 272 (invertebrates) to 72 029 (mammals) parsimony-informative codons. Supplementary Tables 4-6 show the number of informative codons used for each parsimony analysis.

Figure 3 shows the percent overlap of the unrooted trees recovered using the six codon pairing methods (three for parsimony and three for alignment-free) compared to the OTL. For comparison, trees recovered from other alignment-free techniques (CAM, FFP, CVTree, ACS, Andi, and FSWM) and maximum likelihood are also compared to the OTL in Figure 3. Figure 4 shows unrooted tree comparisons of the same algorithms compared to the NCBI taxonomy.

**Figure 1:**
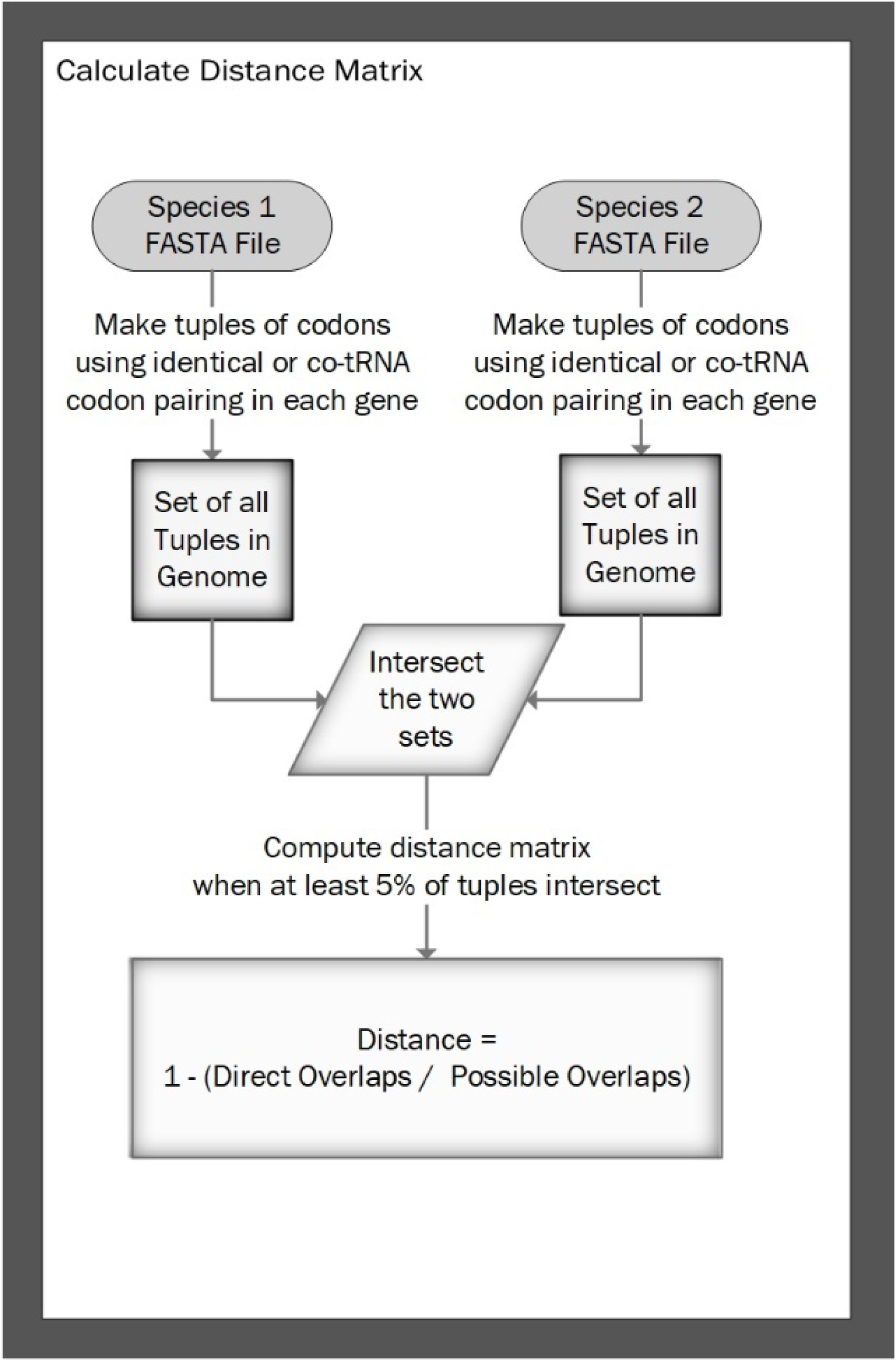
Process to Calculate the Distance Matrix Based on Identical Codon Pairing. Starting with the coding sequences of each gene in a species (FASTA file), codons that use codon pairing within the ribosomal footprint are included in a tuple that is then added to a set for that species. Sets of tuples are intersected to calculate the distance between species. These distances are then added to a distance matrix that can be used to recover phylogenies. Similarly, distances for co-tRNA codon pairing and the combined method are calculated using sets of amino acid tuples instead of codon tuples.

**Figure 2:**
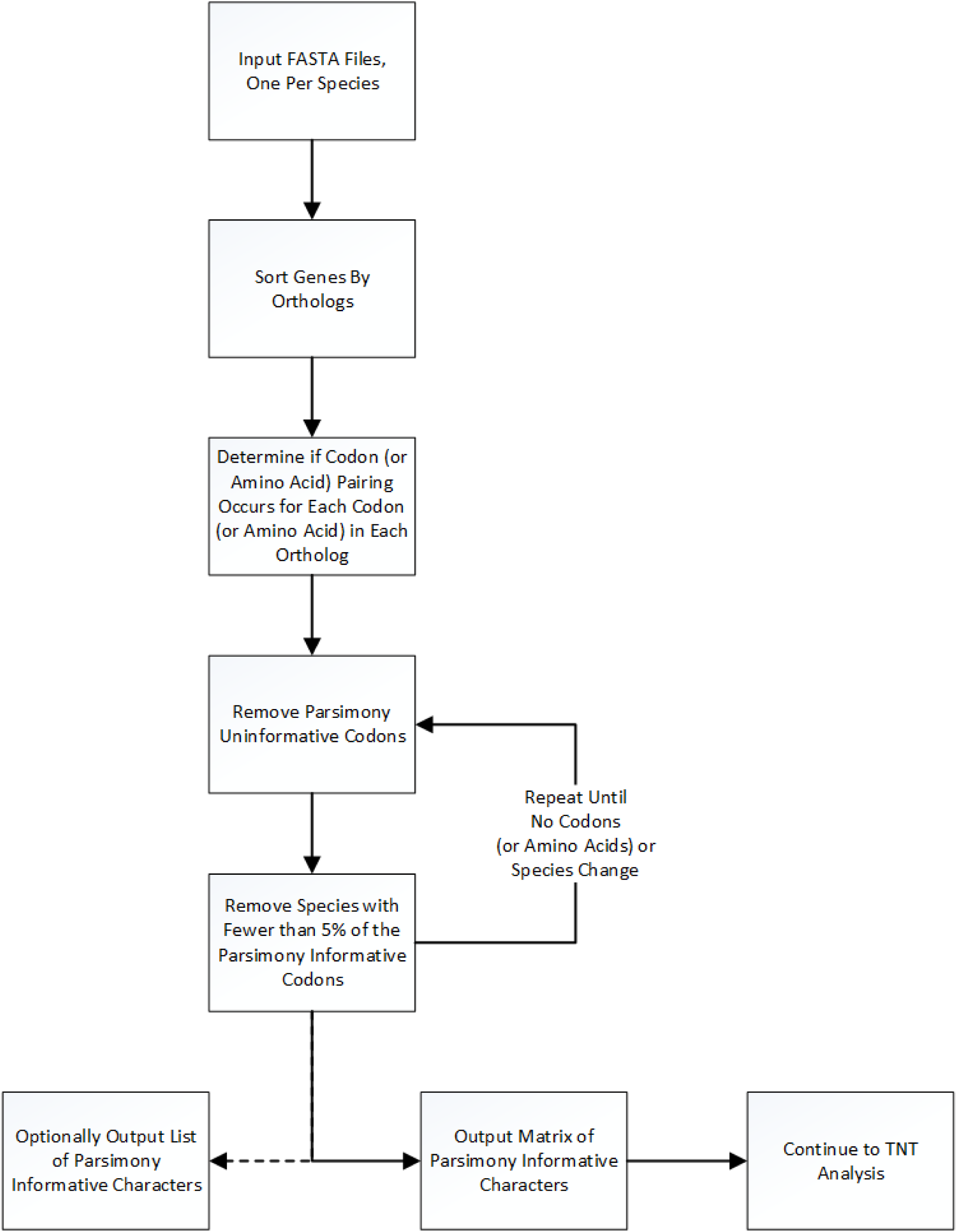
Flow chart for the parsimony analysis. We start with input FASTA files, one for each species. For each codon (or amino acid) within each ortholog, we assign a binary value of ‘0’, ‘1’, or ‘?’ depending on if codon pairing for that codon (or amino acid) occurs. We then remove parsimony-uninformative characters. Next, we remove any species that do not contain at least 5% of the parsimony informative codons, and we conduct the analysis only if at least 5% of the species pass the filter. Finally, we output the parsimony-informative character matrix for each codon (or amino acid) pairing to be used in a TNT analysis and an optional list of parsimony informative characters.

**Figure 3:**
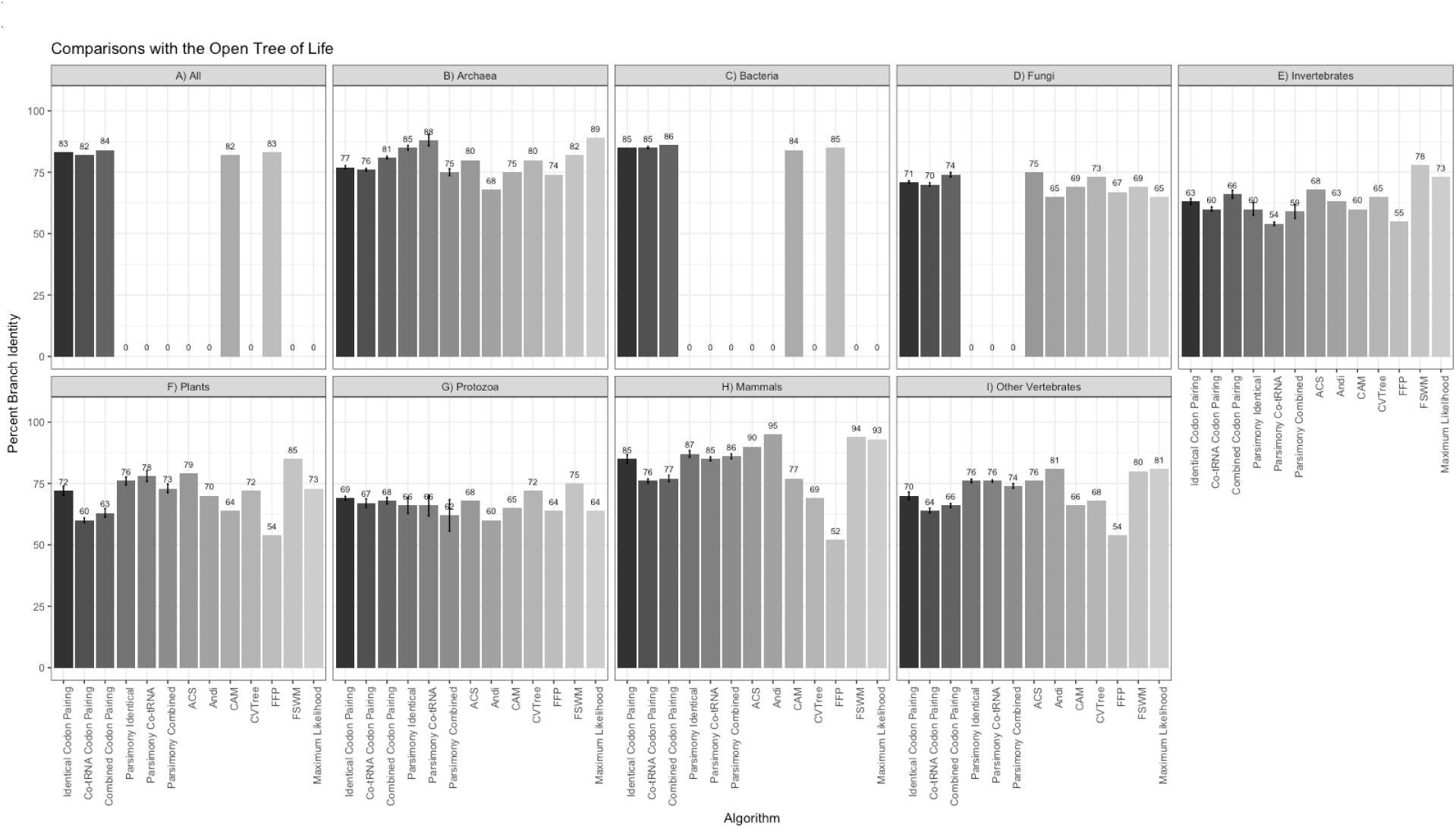
Percent edge overlap for comparisons of each algorithm against the OTL. The alignment-free and parsimony codon pairing methods report the mean percent edge overlap with the OTL based on using different ribosome windows from 2-11. Error bars are reported for the codon pairing methods, signifying one standard deviation from the mean. The other methods were previously reported in Miller, et al. (2019a) and are used for comparison.

**Figure 4:**
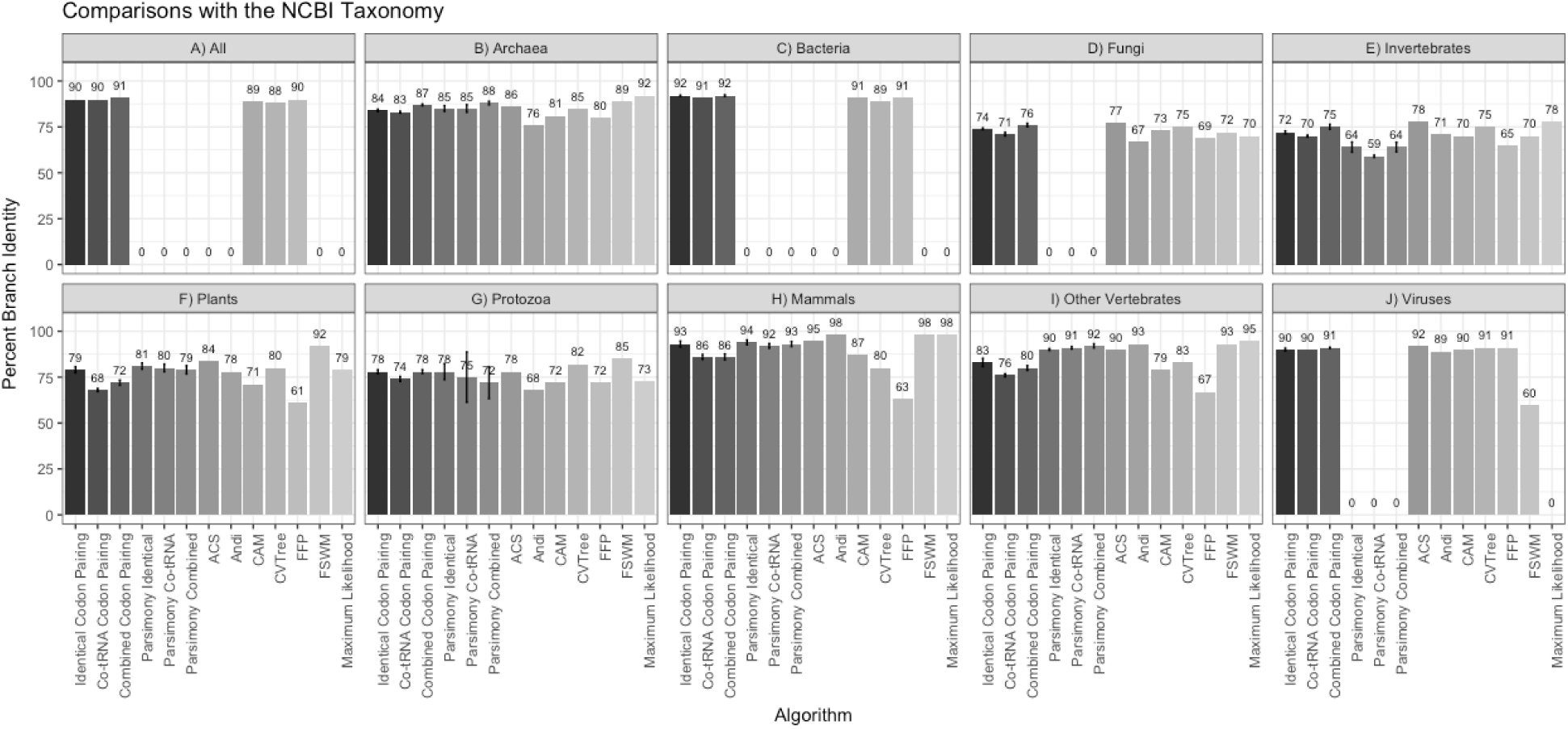
Percent edge overlap for comparisons of each algorithm against the NCBI taxonomy. The alignment-free and parsimony codon pairing methods report the mean percent edge overlap with the NCBI taxonomy based on using different ribosome windows from 2-11. Error bars are reported for the codon pairing methods, signifying one standard deviation from the mean. The other methods were previously reported in Miller, et al. (2019a) and are used for comparison.

The alignment-free codon pairing algorithm and the codon pairing with parsimony recovered phylogenetic relationships that are highly congruent with both the OTL and NCBI database. The alignment-free method had a branch percent identity ranging from 62% −86% with the OTL taxonomy and a range of 68.4% to 92.6% for the NCBI taxonomy. The parsimony pairing methods performed slightly better with branch percent identities ranging from 64% −90% with the OTL and 63.5% to 94% with the NCBI taxonomy. In four of the six taxonomic groups where enough species passed the parsimony filters, the parsimony approach for codon pairing recovered phylogenies that were more congruent with the OTL and the NCBI than the alignment-free approach. Parsimony also tied the alignment-free codon pairing approach in protozoa. The only taxonomic group in which the alignment-free method outperformed parsimony when compared to the OTL was for invertebrates, which also had the fewest parsimony informative characters (See Supplementary Tables 4-6).

The comparisons of the codon pairing algorithms to maximum likelihood and other alignment-free algorithms show that none of the phylogenetic algorithms consistently recovered phylogenies with the highest percent edge similarity with the OTL or the NCBI taxonomy. However, the codon pairing algorithms consistently outperformed CAM and FFP. The codon pairing methods were comparable to maximum likelihood, CVTree, ACS, Andi, and FSWM in all taxonomic groups.

Supplementary Table 7 shows the optimal window sizes and the method (identical, co-tRNA, or combined codon pairing) that recovered the most congruent tree with the established phylogenies. We define the minimum optimal window size as the smallest window size to recover the most congruent phylogeny when compared to the reference. Across all taxonomic groups, the minimum optimal window size was relatively small. Averaged for all minimum optimal window sizes that produced the highest congruence with the OTL, parsimony had a mean minimum optimal window size of 4.000 with a sample standard deviation of 3.033. The alignment-free method had a mean minimum optimal window size of 3.500 with a sample standard deviation of 1.509. Supplementary Tables 8-19 show the percent edge overlap for identical, co-tRNA, and combined codon pairing compared to the OTL and the NCBI taxonomy for both the alignment-free and parsimony approaches at each ribosome window size from 2-11. For both the alignment-free and parsimony approaches, combining co-tRNA codon pairing with identical codon pairing produced the most congruent tree with the OTL and the NCBI Taxonomy in the following taxonomic groups: all species, archaea, bacteria, fungi, invertebrates, protozoa, and viruses. For both methods, identical codon pairing was more congruent with the reference phylogenies in mammals. Parsimony produced a more congruent tree for plants using co-tRNA codon paring, while the alignment-free method had the best results using identical codon pairing. Furthermore, the non-mammalian vertebrate trees were most congruent with the reference phylogenies using identical codon pairing for the alignment-free method and the combined method for parsimony.

We also compared the codon pairing motifs (i.e., the set of codons that paired within a gene) across each taxonomic group. For example, a gene that has identical codon pairing for AAA and AAT would have a motif of {AAA, AAT}. We found that fewer than 10% of codon pairing motifs were identified in multiple species in most taxonomic groups (see Supplementary Figures 1-10). Bacteria had the most repeated codon pairing motifs (13.7%) and fungi had the fewest repeated motifs (0.7%).

The frequency of identical codon pairing was also calculated to determine if systematic biases in pairing based on the codon exist. We counted the number of genes in a species that used identical codon pairing for each codon. We then calculated the frequency of codon pairing for each codon by dividing the number of genes with identical codon pairing for that codon by the total number of genes in that same species. We repeated this process for each codon, creating boxplots of codon pairing frequencies across each taxonomic group (see Supplementary Figures 11-20). Bacteria, archaea, protozoa, and viruses had very wide distributions of codon pairing frequencies. Fungi and invertebrates had narrower distributions of codon pairing frequencies. Mammals, plants, and other vertebrates had very narrow distributions of codon pairing frequencies. Narrow distributions indicate less variability in codon pairing between species among those taxonomic groups. Each taxonomic group has the same pattern of pairing usage (i.e., if a codon pairs frequently in one taxonomic group, it also pairs frequently in other taxonomic groups), although mammals have the least variation between species. Excluding stop codons, codons encoding arginine are the least likely to pair (occurs in ∼20-25% of genes) and codons encoding asparagine and leucine are the most likely to pair (occurs in ∼60-75% of genes), except leucine-encoding CTA, which pairs in ∼20-25% of genes.

We further analyzed the number of codons that paired within each gene. We counted the number of codon pairing motifs that included 1, 2, 3,…, 61 codons and report the distribution for each taxonomic group in Supplementary Figures 21-30. In most taxonomic groups, each motif contains ∼10-40 codons. However, bacteria, archaea, and viruses are more likely to have fewer codons in each motif, while vertebrates typically have more codons in each motif.

Finally, we quantified the frequency of repeated motifs. We counted the number of times each motif was used in each taxonomic group. Supplementary Figures 31-40 show the distribution of repeated motif frequencies in each taxonomic group. In most taxonomic groups, most repeated motifs are repeated 1-20 times with a steep decreasing slope as the motif is repeated more frequently. However, in archaea, the number of times a motif repeats quickly decreases between 1-30 and then the slope increases until 61 before sharply dropping to near zero. The scripts we used to create each supplementary figure can be found at https://github.com/ridgelab/codon_pairing/supplementary_graphs.

## Discussion

Through our analyses, we show that both identical and co-tRNA codon pairing are phylogenetically conserved across all domains of life. We further illustrate that combining identical and co-tRNA codon pairing improves the concordance of recovered phylogenies with the NCBI taxonomy and the OTL in the following taxonomic groups: all species, archaea, bacteria, fungi, invertebrates, protozoa, and viruses. Using parsimony, we also show that combining identical and co-tRNA codon pairing improves the overall concordance of the tree containing non-mammalian vertebrates. In mammals, identical codon pairing had the strongest phylogenetic signal. The most congruent phylogenies for plants were split between only identical codon pairing using the alignment-free method and only co-tRNA codon pairing for the parsimony approach.

This comprehensive analysis shows that codon pairing is a novel class of phylogenetic characters which can provide additional insight into the phylogeny and evolution of organisms. We also provide tools for quickly analyzing thousands of species using our provided framework. As opposed to common ortholog-based techniques that use shared character states to infer phylogenies, identical and co-tRNA codon pairing analyze sequence features that are associated with gene expression. Since gene expression plays a crucial role in adaptive divergence and ecological speciation (Pavey, et al. 2010), and codon pairing affects gene expression, we propose that detectable patterns in codon pairing not only inform phylogeny, but may play a greater role in the speciation process than previously acknowledged. In our analyses, we show that codon pairing alone can recover phylogenies that are comparable to other alignment-free or maximum likelihood approaches.

Our analysis of identical codon pairing found several instances of increased (or decreased) codon pairing within certain codons and amino acids. In some instances, codon pairing (or lack of codon pairing) might be due to protein structure instead of translational efficiency. Arginine (Arg) is very positively charged and highly repulsive to other like-charged amino acids. Although rarely pairing compared to other amino acid residues, arginine pairing is essential to some protein-protein interactions and occurs more frequently than expected by random chance (Lee, et al. 2013). In protein folding, coiled-coil interfaces often make asparagine (Asn)-Asn conformations that face away from the hydrophobic core (Thomas, et al. 2017). Our analysis of codon pairing confirms that asparagine pairing occurs much more frequently than arginine pairing. These interactions suggest that asparagine and arginine pairing conservation might be based on structure instead of codon translational efficiency. In contrast, leucine zipper T cell receptors have the highest expression values (Foley, et al. 2017). Furthermore, the leucine zipper is a 60-80 amino acid protein domain that allows for faster gene expression, sequence-specific DNA-binding, and dimerization (Ellenberger 1994). Our results show that leucin-encoding codons are among the most commonly paired codons. However, leucine-encoding CTA pairs significantly less frequently than other leucine-encoding codons. Further exploration into CTA interactions with other leucine-encoding codons may help determine why CTA pairs much less frequently.

Although co-tRNA codon pairing is less prominent in prokaryotes than in eukaryotes (Shao, et al. 2012; Zhang, et al. 2013; Quax, et al. 2015), we show that identical codon pairing and co-tRNA codon pairing are both phylogenetically conserved in all domains of life. However, we also show that using an alignment-free framework, the most congruent vertebrate and plant phylogenies are generally recovered using only identical codon pairing. Similarly, the parsimony method recovered the most congruent mammal phylogeny using only identical codon pairing. However, parsimony used only co-tRNA codon pairing in plants and the combined approach in non-mammalian vertebrates. We show that although identical and co-tRNA codon pairing do not occur in equal frequencies, they are both phylogenetically conserved. We also show that combining identical and co-tRNA codon pairing recovers phylogenies that most support established phylogenies in seven out of ten taxonomic groups.

Since many orthologous genes are not currently annotated, our alignment-free approach allows researchers to quickly determine where new genomes fit on the OTL without first verifying orthology. In taxonomic groups that include many recently-sequenced genomes, such as bacteria, fungi, and viruses, the alignment-free approach can provide an accurate method to quickly determine the taxonomic relationships of those species. Furthermore, vastly divergent species can be analyzed with a single command at runtime, facilitating the analysis of thousands of species across various taxonomic groups.

In taxonomic groups that have well-documented orthologous relationships, we show that codon pairing recovers parsimony trees that are largely congruent with the OTL and the NCBI taxonomy. Since maximum likelihood has been widely used to establish the reference phylogenies that we used, it is unsurprising that in the most established taxonomic groups, such as mammals and other vertebrates, maximum likelihood recovers trees that are most congruent with the references. However, in plants and protozoa, the parsimony analysis elucidates a phylogenetic signal using only codon pairing that is sufficient to recover more congruent trees with the OTL and the NCBI taxonomy than maximum likelihood. Given the high degree of congruence between the established phylogenies, phylogenies recovered using other techniques, and the trees recovered using only codon pairing, we propose that codon pairing should be considered in future phylogenomic analyses.

## Materials and Methods

### Data Collection and Processing

We downloaded all reference genomes and annotations from the National Center for Biotechnology Information (NCBI) (Pruitt, et al. 2000; Wheeler, et al. 2007; Pruitt, et al. 2014) in September, 2017. Reference genomes were used because they represent the most commonly accepted nucleotides in each species(Pruitt, et al. 2000; Wheeler, et al. 2007). We used the coding sequences (CDS) from the longest isoform of each gene, and we removed genes with previously-annotated exceptions (i.e., translational exception, unclassified transcription discrepancy, suspected errors, partial genes, etc.). A total of 23 423 species were divided into the following taxonomic groups based on NCBI annotations, with some overlap between bacteria and viruses: 418 archaea, 15 063 bacteria, 234 fungi, 149 invertebrates, 89 plants, 75 protozoa, 107 mammalian vertebrates, 123 other vertebrates, and 7 233 viruses. While some of these taxonomic groups, such as invertebrates, do not represent monophyletic clades, we opted to maintain these species classifications to facilitate analyses between different studies that use the NCBI annotations. Furthermore, since our analysis compares different algorithms, all algorithms will be subject to the same potential biases associated with analyzing groups that are not monophyletic.

### Accounting for Differences in Ribosomal Footprint

Estimates of the ribosome footprint vary drastically and can range from 15 nucleotides (5 codons) to about 45 nucleotides (15 codons) with a commonly accepted length of 28 nucleotides (about nine codons) (Martens, et al. 2015). Since codon pairing requires at least two codons, we examined pairing lengths (i.e., a sliding window) of 2-11 codons. This technique allows for variations in the ribosomal footprint among different taxonomic groups and can determine if codon pairing is dispersed throughout the ribosomal footprint or is more phylogenetically conserved at a smaller window size.

### Calculating Identical and co-tRNA Codon Pairing

For both the parsimony and alignment-free methods, we encoded identical codon pairings, co-tRNA codon pairings, and either identical and co-tRNA codon pairings with a binary representation (i.e., if a codon paired within a gene, it was given a value of ‘1’ regardless of the number of times the pairing occurred). We determined which codons used identical codon pairing for each gene by adding each codon that occurred multiple times within the sliding window to a set of codons for that gene. Similarly, we created a set of amino acids for co-tRNA codon pairings for each gene by adding the amino acid product of the paired non-identical codons that encode that amino acid to the ordered set. Since the combined approach considers if either identical or co-tRNA codon pairing occurs, we calculated combined pairing by translating the gene sequence and identifying amino acids that paired within the ribosome window, adding each residue occurring multiple times in the sliding window to a set.

### Alignment-free Codon Pairing Calculation

We present three alignment-free methods to calculate a distance matrix: 1) based on identical codon pairing, 2) based on co-tRNA codon pairing, and 3) based on a combination of either identical or co-tRNA codon pairing. Although genes must be assembled, orthologous relationships are not required or used in the distance matrix calculation. All three methods use a binary (occurs or does not occur) representation of codon pairing within a gene. First, if identical codon pairing occurs anywhere within a gene, the codons are added to a set for that gene. If co-tRNA codon pairing or the combined approach is selected, then amino acids are added to a set if they occur two or more times within the ribosomal footprint anywhere in the gene. Next, the sets are alphabetized and converted to a tuple (immutable list) so they can be added to a set for the entire species. This process is repeated for each gene within a species until all gene pairings have been made into tuples and added to a set for the species. We repeat this process for each species until all species have a set of tuples representing the codons (or amino acids) that are pairing within at least one gene. Finally, we calculate the distances between each species in a pairwise manner. This process is depicted in Figure 1.

Similar to the method used by Miller, et al. (2019a), the pairwise distance between two species, *A* and *B*, is calculated as one minus the relative similarity of the species. The relative similarity of the species is the number of overlapping tuples between the sets of tuples, *a* and *b*, from both species divided by the total number of tuples from *a* or *b* with the fewest number of tuples. This distance is given in equation 1:

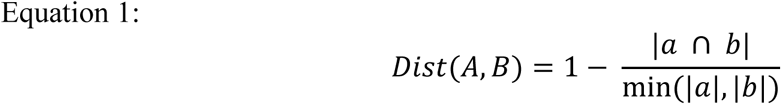

If the ratio of tuples in *a* and *b* does not exceed 5%, the species are assigned the maximum distance of 1.0. This filter limits small genome bias (e.g., without this cutoff, if one gene from a virus with two genes has the same codon pairing profile as a gene in a vertebrate with 20 000 genes, then the distance between the virus and the vertebrate would be 0.5). This process allows us to calculate a distance, with a maximum of 1.0, where more closely related species have a smaller distance because their overall codon pairing biases across all genes are more similar.

### Summary of Alignment-free Options

We implemented `pairing_distance.py` in Python 3.5 to calculate the distance matrix based on the codon pairing algorithm outlined above. We provide several additional options for `pairing_distance.py` to give users greater flexibility in their research. Input FASTA files can be provided either as a list (standard bash expansion) with the `-i` option, or included in a single directory with the `-id` option. The program automatically handles gzipped compressed files with the `.gz` or `.gzip` file extension or uncompressed data with any other file extension. The output distance matrix by default is written to standard out, although an output file can be provided through the `-o` option. Although all available processing cores are used by default to calculate the distance, this can be modified with the `-t` option. RNA sequences can also be provided using the `-rna` flag. The `-l` option allows the user to specify an alternative codon table, with the standard codon table being used by default. By default, the ribosome footprint is set to nine codons, although this option can be modified using `-f`. In the same program, we also provide a flag, `-c`, to allow users to use co-tRNA codon pairing instead of identical codon pairing and the `-b` flag to signify both identical and co-tRNA codon pairing. These options are explained in more detail in the accompanying README file found in the GitHub repository: https://github.com/ridgelab/codon_pairing/tree/master/alignment_free.

### Parsimony Analysis

For our parsimony analysis, we used NCBI ortholog annotations, which include orthologs from a variety of sources. We use Python 3.5 to implement `parsimony_pairing.py` to create a character matrix of parsimony-informative codon pairings from a directory of FASTA files containing gene sequences for each species, one file per species. Each row in the matrix contains a record for a different species. Each column in the matrix represents a parsimony-informative codon (or amino acid) within a specific ortholog. For each species, each codon (or amino acid) in each ortholog is labelled ‘0’ if it does not pair within a ribosomal window, ‘1’ if it does pair, or ‘?’ if the ortholog annotation is unavailable for that species.

To be considered parsimony informative, each included ortholog was present in at least four species, each codon (or amino acid) paired in at least one species, and each codon (or amino acid) did not pair in at least one species. We further required all species to contain at least 5% of all the parsimony-informative codons (or amino acids) to limit the effect of missing data. We created this character matrix and a key file containing an ordered list of each parsimony-informative codon (or amino acid) that was included in the matrix in a single step at runtime (see Figure 2). The following command demonstrates typical usage for identical codon pairing, where `${DIR}` is the path to a directory containing one FASTA file per species, `${MATRIX}` is the path to the output matrix, and `${KEYS}` is the path to the output key file containing the ordered list of parsimony-informative codons.

~~~
python getPairingMatrix.py -id ${DIR} -o ${MATRIX} -oc ${KEYS}
~~~

### Summary of Parsimony Options

We provide the same options in parsimony_pairing.py as the alignment-free method, with a few notable exceptions. In addition to the options described in the alignment-free section, –oc optionally indicates the path to an output file containing the ordered parsimony-informative codons included in the character matrix. Optionally, –on will use a numbering system to create names for the species instead of using the names of the input files. This option is most useful when file names are very long or do not correlate to the species names.

### Constructing Phylogenetic Trees Using Parsimony

We used Tree Analysis Using New Technology (TNT) (Goloboff, et al. 2005) to recover phylogenetic trees using parsimony. We selected TNT based on its ability to handle large datasets and its fast tree-searching algorithms. We found up to 100 most parsimonious trees, saving multiple trees recovered using tree bisection reconnection (tbr) branch swapping (Kumar, et al. 2018).

### Reference Phylogenies

We inferred subtrees of each taxonomic group from both the OTL and the NCBI Taxonomy Browser for each taxonomic group. The OTL combines phylogenetic relationships reported in primary literature and contains a web application programming interface (API) that allows for querying the OTL database. Although the NCBI Taxonomy Browser gathers information from a variety of sources and is therefore not considered a primary source for taxonomic relationships, it contains more species than the OTL and provides added insights into our analyses. We use both phylogenies as reference trees to compare the alignment-free and parsimony trees obtained from codon pairing.

### Open Tree of Life

We used `getOTLtree.py` (Miller, et al. 2019a) to obtain reference trees for each taxonomic group from the OTL in a single step at runtime. This program utilizes the OTL application programming interface (API) to programmatically query the OTL database to first obtain OTL taxonomy identifiers (OTT ids) for each species and then query the OTL database to retrieve the reference tree for the species found. The program also allows users to select the correct domain of life when multiple OTT ids are found for a species (e.g., *Nannospalax galili* is currently listed in the OTL database as both a eukaryote and a bacterium). The output file contains the inferred reference tree from the OTL and a list of any species that the OTL did not include in the tree. We ran this program using the following command, where `${INPUT}` is a list of species, and `${OUTPUT}` is the output file:

~~~
python getOTLtree.py -i ${INPUT} -o ${OUTPUT}
~~~

### NCBI Taxonomy Browser

We used the NCBI taxonomy browser (https://www.ncbi.nlm.nih.gov/Taxonomy/CommonTree/wwwcmt.cgi) to download the taxonomical relationships in PHYLIP (Felsenstein 1989) format. We included unranked taxa to maximize the number of included species for each taxonomic group.

### Tree Comparison

We assessed the accuracy of our identical, co-tRNA, and combined codon pairing algorithms by comparing the trees we recovered to the reference trees from the OTL and the NCBI taxonomy. We determined the similarity between trees by using the ete-compare module from the Environment for Tree Exploration toolkit (ETE3) (Huerta-Cepas, et al. 2016), which computes the percentage of branch similarity between two trees. A higher percentage of branch similarity indicates higher congruence between trees. The branch similarity method has a relatively low computational cost for large datasets, and it allows for unrooted tree comparisons and comparisons of trees with polytomies. For the parsimony analysis, if any taxonomic comparison produced more than one equally parsimonious tree, we computed the percentage of edge similarity between each generated tree and the reference tree. We then reported the average percent overlap of all comparisons.

### Comparison with Maximum Likelihood

We used the same maximum likelihood validation results as previously reported in Miller, et al. (2019a). The ortholog-based maximum likelihood technique first compiled all NCBI ortholog annotations and subsampled the most commonly used orthologs in each taxonomic group, where all gene annotations must be unique within a given species. Next, they used Clustal Omega (Sievers and Higgins 2018) to perform a multiple sequence alignment (MSA) on each orthologous gene cluster. Finally, IQ-TREE (Nguyen, et al. 2015) was used to perform a maximum likelihood analysis on the combined MSA super-matrix from all orthologs. ETE3 was used to compare the recovered phylogenies to the OTL and the NCBI taxonomy. Miller, et al. (2019a) excluded bacteria and viruses from their analyses because of the lack of orthologs spanning a sufficient number of species.

### Comparison with Codon Aversion Motifs

Codon aversion motifs (CAM) are sets of codons that are not used within genes (Miller, et al. 2019a). They have also been used to recover phylogenies using alignment-free techniques. Since our method using codon pairing is also a codon-based method, we included CAM in our comparisons to determine if the phylogenetic signal is more congruent with established phylogenies using codon use/aversion or codon pairing. We use the results reported in Miller, et al. (2019a) for our comparisons.

### Comparison with Feature Frequency Profiles

Comparisons were also done with a k-mer based alignment-free phylogenomic approach, Feature Frequency Profiles (FFP) (Sims, et al. 2009; Jun, et al. 2010). The FFP method works by counting shared k-mers between species, with more directly overlapping k-mer counts being associated with closer species relatedness. Since we use the same dataset as previously reported in Miller, et al. (2019a), we also use their FFP validation set to compare the congruence of FFP with the OTL and the NCBI taxonomy.

### Comparison with CVTree

CVTree is an alignment-free approach that uses composition vectors to calculate the frequency of words of a given length. The algorithm normalizes the composition vector frequencies by expected frequencies from random chance. Species with similar word frequencies would be considered to be closely related. We use the CVTree validation set previously reported in Miller, et al. (2019a).

### Comparison with Average Common Substring (ACS)

ACS is an alignment-free approach that determines distances by calculating the average match lengths of substrings. The algorithm finds the longest substring at each index of a sequence that is also contained in a second sequence. It then calculates the average length of the matching substrings. We use the ACS validation set reported in Miller, et al. (2019a).

### Comparison with andi

Andi is another alignment-free algorithm that searches two genes for areas of exact matches, referred to as anchors, that enclose mismatch areas. It then compares the mismatch areas to compute a distance between species. We use the andi validation set reported in Miller, et al. (2019a).

### Comparison with Filtered Spaced-word Matches

Filtered Spaced-word Matches (FSWM) is an alignment-free approach that is similar to andi because it also searches for matching-spaced words between sequences. FSWM differs from andi by filtering out matches that could have been caused by random chance. We used the FSWM validation set reported in Miller, et al. (2019a).

## Acknowledgements

We appreciate the support of Brigham Young University and the technical assistance of the Fulton Supercomputing Laboratory staff.

