## Supplemental Material for "Codon Pairs are Phylogenetically Conserved: Codon pairing as a new class of phylogenetic characters"

Supplementary Information

### Tables

Supplementary Table 1: Species Included in Identical Codon Pairing

| Taxonomic Group | 2 | 3 | 4 | 5 | 6 | 7 | 8 | 9 | 10 | 11 | Average | Total Number of Species |
|---|---|---|---|---|---|---|---|---|---|---|---|---|
| **Archaea** | 106 | 95 | 95 | 95 | 95 | 95 | 95 | 95 | 95 | 95 | 96.1 | 418 |
| **Fungi** | 19 | 9 | 13 | 9 | 9 | 19 | 9 | 13 | 9 | 9 | 11.8 | 234 |
| **Invertebrates** | 65 | 55 | 57 | 55 | 55 | 55 | 63 | 55 | 57 | 55 | 57.2 | 149 |
| **Plants** | 60 | 60 | 60 | 61 | 61 | 61 | 61 | 61 | 61 | 61 | 60.7 | 89 |
| **Protozoa** | 15 | 15 | 15 | 16 | 17 | 16 | 16 | 20 | 15 | 15 | 16 | 75 |
| **Mammals** | 97 | 97 | 97 | 97 | 97 | 97 | 97 | 97 | 97 | 97 | 97 | 107 |
| **Other vertebrates** | 114 | 114 | 114 | 114 | 114 | 114 | 114 | 114 | 114 | 114 | 114 | 123 |
| **Viruses** | 168 | 137 | 152 | 188 | 177 | 174 | 174 | 220 | 176 | 184 | 175 | 7 233 |

*Each column is the length of the ribosome window, in codons, that was used and each cell is the number of species that included at least 5% of the parsimony-informative codons for each taxonomic group. Column 12 is the average number of species included in each ribosome window. Column 13 is the total number of species in each taxonomic group.*

Supplementary Table 2: Species Used in Co-tRNA Codon Pairing

| Taxonomic Group | 2 | 3 | 4 | 5 | 6 | 7 | 8 | 9 | 10 | 11 | Average | Total Number of Species |
|---|---|---|---|---|---|---|---|---|---|---|---|---|
| **Archaea** | 107 | 107 | 107 | 107 | 107 | 107 | 107 | 107 | 107 | 106 | 106.9 | 418 |
| **Fungi** | 9 | 10 | 10 | 20 | 9 | 20 | 9 | 14 | 13 | 10 | 12.7 | 234 |
| **Invertebrates** | 65 | 65 | 66 | 55 | 65 | 57 | 65 | 55 | 57 | 55 | 60.5 | 149 |
| **Plants** | 61 | 59 | 59 | 59 | 59 | 59 | 59 | 59 | 59 | 59 | 59.2 | 89 |
| **Protozoa** | 17 | 16 | 19 | 17 | 15 | 18 | 15 | 16 | 16 | 16 | 16.5 | 75 |
| **Mammals** | 97 | 97 | 97 | 97 | 97 | 97 | 97 | 97 | 97 | 97 | 97 | 107 |
| **Other vertebrates** | 114 | 114 | 114 | 114 | 114 | 114 | 114 | 114 | 114 | 114 | 114 | 123 |
| **Viruses** | 282 | 279 | 262 | 247 | 243 | 256 | 256 | 245 | 236 | 244 | 255 | 7 233 |

*Each column is the length of the ribosome window, in codons, that was used and each cell is the number of species that included at least 5% of the parsimony-informative amino acids for each taxonomic group. Column 12 is the average number of species included in each ribosome window. Column 13 is the total number of species in each taxonomic group.*

Supplementary Table 3: Species Used in Combined Comparison

| Taxonomic Group | 2 | 3 | 4 | 5 | 6 | 7 | 8 | 9 | 10 | 11 | Average | Total Number of Species |
|---|---|---|---|---|---|---|---|---|---|---|---|---|
| Archaea | 96 | 100 | 96 | 100 | 101 | 101 | 101 | 95 | 105 | 106 | 100.1 | 418 |
| Fungi | 13 | 13 | 10 | 14 | 11 | 11 | 13 | 10 | 11 | 14 | 12 | 234 |
| Invertebrates | 55 | 57 | 57 | 69 | 93 | 58 | 57 | 65 | 65 | 58 | 63.4 | 149 |
| Plants | 61 | 61 | 59 | 59 | 59 | 59 | 59 | 59 | 59 | 59 | 59.4 | 89 |
| Protozoa | 15 | 15 | 25 | 27 | 17 | 18 | 21 | 20 | 20 | 21 | 19.9 | 75 |
| Mammals | 97 | 97 | 97 | 97 | 97 | 97 | 97 | 97 | 97 | 97 | 97 | 107 |
| Other vertebrates | 114 | 114 | 114 | 114 | 114 | 114 | 114 | 114 | 114 | 114 | 114 | 123 |
| Viruses | 199 | 224 | 180 | 190 | 190 | 180 | 278 | 262 | 261 | 257 | 222.1 | 7 233 |

*Each column is the length of the ribosome window, in codons, that was used and each cell is the number of species that included at least 5% of the parsimony-informative amino acids for each taxonomic group. Column 12 is the average number of species included in each ribosome window. Column 13 is the total number of species in each taxonomic group.*

Supplementary Table 4: Parsimony Informative Codons used in Identical Codon Pairing

| Taxonomic Group | 2 | 3 | 4 | 5 | 6 | 7 | 8 | 9 | 10 | 11 | Average |
|---|---|---|---|---|---|---|---|---|---|---|---|
| Archaea | 6151 | 8450 | 9902 | 10544 | 11035 | 11254 | 11518 | 11687 | 11664 | 11842 | 10404.7 |
| Fungi | N/A | N/A | N/A | N/A | N/A | N/A | N/A | N/A | N/A | N/A | N/A |
| Invertebrates | 794 | 988 | 1081 | 1160 | 1236 | 1263 | 1329 | 1427 | 1353 | 1423 | 1205.4 |
| Plants | 6230 | 8033 | 9036 | 9842 | 10153 | 10517 | 10607 | 10691 | 10725 | 10693 | 9652.7 |
| Protozoa | 12449 | 14864 | 16051 | 16253 | 16103 | 16171 | 15837 | 15838 | 15764 | 15532 | 15486.2 |
| Mammals | 197074 | 311796 | 319490 | 381908 | 404078 | 335058 | 398896 | 386949 | 436474 | 380879 | 355260.2 |
| Other vertebrates | 228194 | 277024 | 347408 | 388789 | 355121 | 376798 | 400771 | 380813 | 390161 | 15532 | 316061.1 |
| Viruses | 16622 | 23528 | 28145 | 28776 | 30176 | 30768 | 32248 | 31580 | 33374 | 33082 | 28829.9 |

Supplementary Table 5: Parsimony Informative Codons used in Co-tRNA Codon Pairing

| Taxonomic Group | 2 | 3 | 4 | 5 | 6 | 7 | 8 | 9 | 10 | 11 | Average |
|---|---|---|---|---|---|---|---|---|---|---|---|
| Archaea | 3293 | 3294 | 3087 | 2921 | 2783 | 2725 | 2689 | 2661 | 2560 | 2579 | 2859.2 |
| Fungi | N/A | N/A | N/A | N/A | N/A | N/A | N/A | N/A | N/A | N/A | N/A |
| Invertebrates | 418 | 461 | 475 | 455 | 450 | 429 | 422 | 428 | 410 | 382 | 433 |
| Plants | 3219 | 3188 | 3157 | 3082 | 2945 | 2940 | 2929 | 2819 | 2808 | 2723 | 2981 |
| Protozoa | 5415 | 5319 | 5020 | 4840 | 4589 | 4358 | 4198 | 4067 | 3976 | 2819 | 4460.1 |
| Mammals | 94018 | 10195 | 93587 | 93627 | 82729 | 74208 | 70805 | 71310 | 69678 | 55666 | 71582.3 |
| Other vertebrates | 90872 | 86618 | 82126 | 73534 | 74286 | 70704 | 63995 | 62945 | 60569 | 57241 | 72289 |
| Viruses | 11409 | 12103 | 11556 | 11421 | 11248 | 11068 | 10812 | 10550 | 10325 | 10115 | 11060.7 |

Supplementary Table 6: Parsimony Informative Codons used in the Combined Analysis

| Taxonomic Group | 2 | 3 | 4 | 5 | 6 | 7 | 8 | 9 | 10 | 11 | Average |
|---|---|---|---|---|---|---|---|---|---|---|---|
| **Archaea** | 2823 | 2568 | 2292 | 2105 | 1974 | 1851 | 1756 | 1577 | 1612 | 1527 | 2008.5 |
| **Fungi** | N/A | N/A | N/A | N/A | N/A | N/A | N/A | N/A | N/A | N/A | N/A |
| **Invertebrates** | 463 | 464 | 417 | 411 | 359 | 351 | 319 | 302 | 293 | 272 | 365.1 |
| **Plants** | 3236 | 2813 | 2571 | 2347 | 2214 | 2085 | 1977 | 1849 | 1789 | 1709 | 2259 |
| **Protozoa** | 4488 | 3603 | 3021 | 2813 | 2381 | 2311 | 2122 | 1951 | 1862 | 1730 | 2628.2 |
| **Mammals** | 72029 | 64087 | 56125 | 44448 | 44984 | 37830 | 45077 | 41319 | 34711 | 35801 | 44089.2 |
| **Other vertebrates** | 6937 | 61754 | 48110 | 45789 | 43517 | 41519 | 34718 | 33856 | 30895 | 27919 | 37501.4 |
| **Viruses** | 12737 | 12003 | 11587 | 10946 | 10352 | 9924 | 9691 | 9247 | 8921 | 8578 | 10398.6 |

Supplementary Table 7: Optimal Window Size and Options for Each Taxonomic Group

| Taxonomic Group | Alignment-free identical codon pairing (I), co-tRNA codon pairing (C), or both (B) | Alignment-free minimum optimal window sizes | Maximum parsimony identical codon pairing (I), co-tRNA codon pairing (C), or both (B) | Maximum parsimony minimum optimal window sizes |
|---|---|---|---|---|
| All | B | 2 | N/A | N/A |
| Archaea | B | 4 | B | 3 |
| Bacteria* | B | 2 | N/A | N/A |
| Fungi | B | 5 | N/A | N/A |
| Invertebrates | B | 2 | B | 4 |
| Plants | I | 4 | C | 10 |
| Protozoa | B | 2 | B | 2 |
| Mammals | I | 6 | I | 2 |
| Other vertebrates | I | 5 | B | 3 |
| Viruses* | B | 3 | N/A | N/A |

*The first column indicates the taxonomic group. Column 2 shows if the best percent overlap for the alignment-free method came from identical codon pairing (I), co-tRNA codon pairing (C), or the combined method (B). Column 3 shows the window size used to recover the phylogeny most congruent with the NCBI Taxonomy and the OTL for the alignment-free method. If multiple window sizes recovered phylogenies with the same percent congruence, preference was given to smaller window sizes. Column 4 shows if the best percent overlap for the parsimony method came from identical codon pairing (I), co-tRNA codon pairing (C), or the combined method (B). Column 5 shows the window size used to recover the phylogeny most congruent with the NCBI Taxonomy and the OTL for the parsimony method. If multiple window sizes recovered phylogenies with the same percent congruence, preference was given to smaller window sizes. *Indicates that some species overlap between the viruses and bacteria.*

Supplementary Table 8:

| Taxonomic Group | 2 | 3 | 4 | 5 | 6 | 7 | 8 | 9 | 10 | 11 |
|---|---|---|---|---|---|---|---|---|---|---|
| All | 90 | 90 | 90 | 90 | 90 | 90 | 90 | 90 | 90 | 90 |
| Archaea | 84 | 84 | 84 | 84 | 85 | 84 | 84 | 83 | 83 | 83 |
| Bacteria | 91 | 92 | 92 | 92 | 92 | 92 | 92 | 92 | 92 | 92 |
| Fungi | 74 | 73 | 74 | 74 | 74 | 75 | 75 | 74 | 74 | 74 |
| Invertebrates | 72 | 72 | 72 | 72 | 73 | 74 | 72 | 72 | 72 | 71 |
| Plants | 75 | 78 | 81 | 80 | 80 | 79 | 80 | 80 | 80 | 80 |
| Protozoa | 79 | 78 | 79 | 79 | 77 | 78 | 76 | 77 | 77 | 77 |
| Mammals | 92 | 92 | 94 | 94 | 95 | 94 | 93 | 91 | 90 | 91 |
| Other Vertebrates | 81 | 85 | 86 | 87 | 85 | 83 | 81 | 82 | 82 | 81 |
| Viruses | 89 | 89 | 89 | 90 | 90 | 91 | 91 | 91 | 90 | 90 |

*Alignment-free: Identical codon pairing percent overlap with the NCBI Taxonomy. Column 1 shows the taxonomic groups analyzed. Columns 2-11 show the percent overlap with the NCBI Taxonomy from a phylogeny recovered using identical codon pairing with the respective window size 2-11. The highest percent overlap for each taxonomic group is highlighted.*

Supplementary Table 9:

| Taxonomic Group | 2 | 3 | 4 | 5 | 6 | 7 | 8 | 9 | 10 | 11 |
|---|---|---|---|---|---|---|---|---|---|---|
| All | 83 | 83 | 83 | 83 | 83 | 83 | 83 | 83 | 83 | 83 |
| Archaea | 78 | 78 | 77 | 78 | 78 | 77 | 77 | 77 | 77 | 76 |
| Bacteria | 85 | 85 | 85 | 85 | 85 | 85 | 85 | 85 | 85 | 85 |
| Fungi | 71 | 72 | 71 | 71 | 71 | 72 | 72 | 71 | 71 | 72 |
| Invertebrates | 63 | 64 | 63 | 63 | 65 | 65 | 64 | 63 | 62 | 61 |
| Plants | 68 | 71 | 73 | 72 | 72 | 71 | 74 | 72 | 74 | 74 |
| Protozoa | 69 | 70 | 70 | 70 | 68 | 70 | 69 | 69 | 69 | 68 |
| Mammals | 83 | 85 | 86 | 86 | 89 | 87 | 86 | 84 | 84 | 84 |
| Other Vertebrates | 68 | 71 | 72 | 73 | 72 | 71 | 69 | 70 | 70 | 69 |

*Alignment-free: Identical codon pairing percent overlap with the OTL. Column 1 shows the taxonomic group analyzed. Columns 2-11 show the percent overlap with the OTL from a phylogeny recovered using identical codon pairing with the respective window size 2-11. The highest percent overlap for each taxonomic group is highlighted.*

Supplementary Table 10:

| Taxonomic Group | 2 | 3 | 4 | 5 | 6 | 7 | 8 | 9 | 10 | 11 |
|---|---|---|---|---|---|---|---|---|---|---|
| All | 91 | 91 | 91 | 91 | 91 | 91 | 91 | 91 | 91 | 91 |
| Archaea | 88 | 87 | 88 | 88 | 87 | 87 | 87 | 87 | 87 | 87 |
| Bacteria | 93 | 93 | 93 | 92 | 92 | 92 | 92 | 92 | 92 | 92 |
| Fungi | 77 | 77 | 77 | 78 | 77 | 77 | 76 | 75 | 76 | 75 |
| Invertebrates | 78 | 76 | 75 | 75 | 75 | 74 | 74 | 74 | 74 | 73 |
| Plants | 74 | 73 | 73 | 73 | 73 | 71 | 72 | 70 | 71 | 70 |
| Protozoa | 80 | 79 | 79 | 78 | 77 | 77 | 77 | 77 | 78 | 78 |
| Mammals | 89 | 87 | 87 | 87 | 85 | 85 | 85 | 85 | 84 | 84 |
| Other Vertebrates | 79 | 78 | 77 | 80 | 81 | 80 | 81 | 81 | 80 | 80 |
| Viruses | 90 | 91 | 91 | 91 | 91 | 91 | 91 | 91 | 91 | 91 |

*Alignment-free: Both co-tRNA and identical codon pairing percent overlap with the NCBI Taxonomy. Column 1 shows the taxonomic group analyzed. Columns 2-11 show the percent overlap with the NCBI Taxonomy from a phylogeny recovered using both co-tRNA and identical codon pairing with the respective window size 2-11. The highest percent overlap for each taxonomic group is highlighted.*

Supplementary Table 11:

| Taxonomic Group | 2 | 3 | 4 | 5 | 6 | 7 | 8 | 9 | 10 | 11 |
|---|---|---|---|---|---|---|---|---|---|---|
| All | 84 | 84 | 84 | 84 | 84 | 84 | 84 | 84 | 84 | 84 |
| Archaea | 81 | 81 | 82 | 81 | 81 | 81 | 81 | 81 | 80 | 81 |
| Bacteria | 86 | 86 | 86 | 86 | 86 | 86 | 86 | 86 | 86 | 86 |
| Fungi | 74 | 75 | 74 | 76 | 74 | 74 | 73 | 73 | 74 | 73 |
| Invertebrates | 69 | 67 | 67 | 65 | 66 | 65 | 64 | 65 | 64 | 64 |
| Plants | 65 | 65 | 65 | 65 | 63 | 62 | 62 | 61 | 62 | 61 |
| Protozoa | 70 | 69 | 70 | 69 | 67 | 67 | 67 | 67 | 67 | 67 |
| Mammals | 79 | 76 | 78 | 78 | 76 | 76 | 76 | 77 | 75 | 75 |
| Other Vertebrates | 67 | 66 | 65 | 66 | 66 | 67 | 67 | 68 | 67 | 66 |

*Alignment-free: Both co-tRNA and identical codon pairing percent overlap with the OTL. Column 1 shows the taxonomic group analyzed. Columns 2-11 show the percent overlap with the OTL from a phylogeny recovered using both identical and co-tRNA codon pairing with the respective window size 2-11. The highest percent overlap for each taxonomic group is highlighted.*

Supplementary Table 12:

| Taxonomic Group | 2 | 3 | 4 | 5 | 6 | 7 | 8 | 9 | 10 | 11 |
|---|---|---|---|---|---|---|---|---|---|---|
| All | 90 | 90 | 90 | 90 | 90 | 90 | 90 | 90 | 90 | 90 |
| Archaea | 83 | 83 | 83 | 83 | 83 | 82 | 82 | 82 | 83 | 82 |
| Bacteria | 91 | 91 | 91 | 91 | 91 | 91 | 91 | 91 | 91 | 91 |
| Fungi | 72 | 72 | 72 | 73 | 72 | 71 | 71 | 70 | 71 | 70 |
| Invertebrates | 70 | 70 | 70 | 70 | 70 | 70 | 69 | 69 | 69 | 69 |
| Plants | 70 | 69 | 69 | 68 | 68 | 68 | 68 | 69 | 67 | 68 |
| Protozoa | 76 | 75 | 77 | 75 | 73 | 74 | 73 | 74 | 73 | 75 |
| Mammals | 87 | 87 | 86 | 87 | 85 | 86 | 87 | 84 | 84 | 84 |
| Other Vertebrates | 76 | 76 | 76 | 76 | 76 | 76 | 75 | 74 | 77 | 75 |
| Viruses | 90 | 90 | 90 | 90 | 90 | 90 | 90 | 90 | 90 | 90 |

*Alignment-free: Co-tRNA codon pairing percent overlap with the NCBI Taxonomy. Column 1 shows the taxonomic group analyzed. Columns 2-11 show the percent overlap with the NCBI Taxonomy from a phylogeny recovered using co-tRNA codon pairing with the respective window size 2-11. The highest percent overlap for each taxonomic group is highlighted.*

Supplementary Table 13:

| Taxonomic Group | 2 | 3 | 4 | 5 | 6 | 7 | 8 | 9 | 10 | 11 |
|---|---|---|---|---|---|---|---|---|---|---|
| All | 82 | 82 | 82 | 82 | 82 | 82 | 82 | 82 | 82 | 82 |
| Archaea | 75 | 76 | 75 | 76 | 76 | 75 | 75 | 75 | 76 | 76 |
| Bacteria | 84 | 84 | 85 | 85 | 85 | 85 | 85 | 85 | 85 | 85 |
| Fungi | 69 | 70 | 71 | 70 | 70 | 69 | 69 | 69 | 69 | 69 |
| Invertebrates | 62 | 61 | 61 | 61 | 60 | 60 | 60 | 61 | 60 | 59 |
| Plants | 62 | 60 | 60 | 60 | 59 | 59 | 59 | 60 | 59 | 59 |
| Protozoa | 70 | 68 | 69 | 68 | 65 | 66 | 65 | 66 | 65 | 66 |
| Mammals | 77 | 77 | 76 | 77 | 76 | 77 | 77 | 76 | 75 | 75 |
| Other Vertebrates | 66 | 65 | 63 | 64 | 64 | 65 | 65 | 64 | 65 | 63 |

*Alignment-free: Co-tRNA codon pairing percent overlap with the OTL. Column 1 shows the taxonomic group analyzed. Columns 2-11 show the percent overlap with the OTL from a phylogeny recovered using co-tRNA codon pairing with the respective window size 2-11. The highest percent overlap for each taxonomic group is highlighted.*

**Parsimony**

Supplementary Table 14:

| Taxonomic Group | 2 | 3 | 4 | 5 | 6 | 7 | 8 | 9 | 10 | 11 |
|---|---|---|---|---|---|---|---|---|---|---|
| All | N/A | N/A | N/A | N/A | N/A | N/A | N/A | N/A | N/A | N/A |
| Archaea | 84 | 83 | 87 | 88 | 86 | 85 | 86 | 84 | 85 | 84 |
| Bacteria | N/A | N/A | N/A | N/A | N/A | N/A | N/A | N/A | N/A | N/A |
| Fungi | N/A | N/A | N/A | N/A | N/A | N/A | N/A | N/A | N/A | N/A |
| Invertebrates | 58 | 63 | 67 | 65 | 67 | 64 | 61 | 65 | 62 | 63 |
| Plants | 80 | 80 | 82 | 82 | 81 | 82 | 84 | 79 | 79 | 79 |
| Protozoa | 81 | 81 | 81 | 77 | 73 | 77 | 77 | 68 | 81 | 81 |
| Mammals | 96 | 96 | 95 | 94 | 93 | 92 | 93 | 94 | 94 | 93 |
| Other Vertebrates | 91 | 91 | 90 | 90 | 91 | 90 | 90 | 91 | 90 | 91 |
| Viruses | N/A | N/A | N/A | N/A | N/A | N/A | N/A | N/A | N/A | N/A |

*Parsimony: Identical codon pairing percent overlap with the NCBI Taxonomy. Column 1 shows the taxonomic group analyzed. Columns 2-11 show the percent overlap with the NCBI Taxonomy from a phylogeny recovered using identical codon pairing with the respective window size 2-11. The highest percent overlap for each taxonomic group is highlighted.*

Supplementary Table 15:

| Taxonomic Group | 2 | 3 | 4 | 5 | 6 | 7 | 8 | 9 | 10 | 11 |
|---|---|---|---|---|---|---|---|---|---|---|
| All | N/A | N/A | N/A | N/A | N/A | N/A | N/A | N/A | N/A | N/A |
| Archaea | 84 | 84 | 86 | 86 | 86 | 86 | 85 | 84 | 85 | 84 |
| Bacteria | N/A | N/A | N/A | N/A | N/A | N/A | N/A | N/A | N/A | N/A |
| Fungi | N/A | N/A | N/A | N/A | N/A | N/A | N/A | N/A | N/A | N/A |
| Invertebrates | 55 | 58 | 62 | 61 | 64 | 59 | 57 | 61 | 59 | 60 |
| Plants | 77 | 75 | 77 | 78 | 78 | 77 | 79 | 75 | 74 | 75 |
| Protozoa | 68 | 68 | 68 | 66 | 63 | 66 | 66 | 58 | 68 | 68 |
| Mammals | 90 | 89 | 87 | 86 | 86 | 86 | 86 | 88 | 87 | 87 |
| Other Vertebrates | 77 | 76 | 75 | 76 | 76 | 76 | 75 | 76 | 77 | 77 |

*Parsimony: Identical codon pairing percent overlap with the OTL. Column 1 shows the taxonomic group analyzed. Columns 2-11 show the percent overlap with the OTL from a phylogeny recovered using identical codon pairing with the respective window size 2-11. The highest percent overlap for each taxonomic group is highlighted.*

Supplementary Table 16:

| Taxonomic Group | 2 | 3 | 4 | 5 | 6 | 7 | 8 | 9 | 10 | 11 |
|---|---|---|---|---|---|---|---|---|---|---|
| **All** | N/A | N/A | N/A | N/A | N/A | N/A | N/A | N/A | N/A | N/A |
| **Archaea** | 87 | 90 | 89 | 89 | 88 | 87 | 88 | 88 | 89 | 87 |
| **Bacteria** | N/A | N/A | N/A | N/A | N/A | N/A | N/A | N/A | N/A | N/A |
| **Fungi** | N/A | N/A | N/A | N/A | N/A | N/A | N/A | N/A | N/A | N/A |
| **Invertebrates** | 63 | 65 | 71 | 63 | 63 | 63 | 63 | 63 | 62 | 64 |
| **Plants** | 77 | 77 | 79 | 77 | 84 | 81 | 78 | 80 | 77 | 78 |
| **Protozoa** | 87 | 85 | 67 | 66 | 76 | 75 | 67 | 67 | 61 | 66 |
| **Mammals** | 95 | 92 | 95 | 91 | 94 | 92 | 92 | 93 | 94 | 94 |
| **Other Vertebrates** | 93 | 94 | 92 | 93 | 93 | 93 | 91 | 91 | 91 | 91 |
| **Viruses** | N/A | N/A | N/A | N/A | N/A | N/A | N/A | N/A | N/A | N/A |

*Parsimony: Combined codon pairing percent overlap with the NCBI Taxonomy. Column 1 shows the taxonomic group analyzed. Columns 2-11 show the percent overlap with the NCBI Taxonomy from a phylogeny recovered using co-tRNA codon pairing with the respective window size 2-11. The highest percent overlap for each taxonomic group is highlighted.*

Supplementary Table 17:

| Taxonomic Group | 2 | 3 | 4 | 5 | 6 | 7 | 8 | 9 | 10 | 11 |
|---|---|---|---|---|---|---|---|---|---|---|
| **All** | N/A | N/A | N/A | N/A | N/A | N/A | N/A | N/A | N/A | N/A |
| **Archaea** | 86 | 87 | 90 | 89 | 89 | 86 | 88 | 88 | 87 | 86 |
| **Bacteria** | N/A | N/A | N/A | N/A | N/A | N/A | N/A | N/A | N/A | N/A |
| **Fungi** | N/A | N/A | N/A | N/A | N/A | N/A | N/A | N/A | N/A | N/A |
| **Invertebrates** | 57 | 61 | 66 | 57 | 56 | 58 | 59 | 59 | 58 | 57 |
| **Plants** | 73 | 71 | 74 | 72 | 77 | 75 | 73 | 74 | 72 | 72 |
| **Protozoa** | 73 | 71 | 57 | 57 | 66 | 66 | 59 | 59 | 54 | 60 |
| **Mammals** | 87 | 86 | 86 | 85 | 87 | 85 | 84 | 85 | 85 | 85 |
| **Other Vertebrates** | N/A | N/A | N/A | N/A | N/A | N/A | N/A | N/A | N/A | N/A |

*Parsimony: Combined codon pairing percent overlap with the OTL. Column 1 shows the taxonomic group analyzed. Columns 2-11 show the percent overlap with the OTL from a phylogeny recovered using co-tRNA codon pairing with the respective window size 2-11. The highest percent overlap for each taxonomic group is highlighted.*

Supplementary Table 18:

| Taxonomic Group | 2 | 3 | 4 | 5 | 6 | 7 | 8 | 9 | 10 | 11 |
|---|---|---|---|---|---|---|---|---|---|---|
| All | N/A | N/A | N/A | N/A | N/A | N/A | N/A | N/A | N/A | N/A |
| Archaea | 88 | 88 | 86 | 86 | 83 | 84 | 84 | 83 | 85 | 82 |
| Bacteria | N/A | N/A | N/A | N/A | N/A | N/A | N/A | N/A | N/A | N/A |
| Fungi | N/A | N/A | N/A | N/A | N/A | N/A | N/A | N/A | N/A | N/A |
| Invertebrates | 59 | 60 | 59 | 60 | 59 | 58 | 59 | 59 | 58 | 59 |
| Plants | 76 | 82 | 80 | 80 | 80 | 82 | 81 | 80 | 84 | 80 |
| Protozoa | 38 | 77 | 74 | 75 | 86 | 74 | 86 | 77 | 81 | 81 |
| Mammals | 94 | 92 | 91 | 92 | 93 | 90 | 94 | 92 | 92 | 92 |
| Other Vertebrates | 92 | 91 | 91 | 91 | 90 | 90 | 91 | 91 | 90 | 90 |
| Viruses | N/A | N/A | N/A | N/A | N/A | N/A | N/A | N/A | N/A | N/A |

*Parsimony: Co-tRNA codon pairing percent overlap with the NCBI Taxonomy. Column 1 shows the taxonomic group analyzed. Columns 2-11 show the percent overlap with the NCBI Taxonomy from a phylogeny recovered using co-tRNA codon pairing with the respective window size 2-11. The highest percent overlap for each taxonomic group is highlighted.*

Supplementary Table 19:

| Taxonomic Group | 2 | 3 | 4 | 5 | 6 | 7 | 8 | 9 | 10 | 11 |
|---|---|---|---|---|---|---|---|---|---|---|
| All | N/A | N/A | N/A | N/A | N/A | N/A | N/A | N/A | N/A | N/A |
| Archaea | 89 | 88 | 86 | 87 | 83 | 85 | 85 | 82 | 83 | 84 |
| Bacteria | N/A | N/A | N/A | N/A | N/A | N/A | N/A | N/A | N/A | N/A |
| Fungi | N/A | N/A | N/A | N/A | N/A | N/A | N/A | N/A | N/A | N/A |
| Invertebrates | 53 | 54 | 54 | 54 | 53 | 55 | 54 | 53 | 53 | 53 |
| Plants | 73 | 75 | 77 | 76 | 79 | 79 | 79 | 79 | 80 | 78 |
| Protozoa | 59 | 66 | 63 | 62 | 72 | 65 | 72 | 66 | 69 | 69 |
| Mammals | 86 | 85 | 85 | 86 | 85 | 84 | 86 | 84 | 85 | 84 |
| Other Vertebrates | 76 | 76 | 77 | 76 | 75 | 76 | 76 | 76 | 76 | 76 |

*Parsimony: Co-tRNA codon pairing percent overlap with the OTL. Column 1 shows the taxonomic group analyzed. Columns 2-11 show the percent overlap with the OTL from a phylogeny recovered using co-tRNA codon pairing with the respective window size 2-11. The highest percent overlap for each taxonomic group is highlighted.*

**Figures**

Supplementary Figure 1:

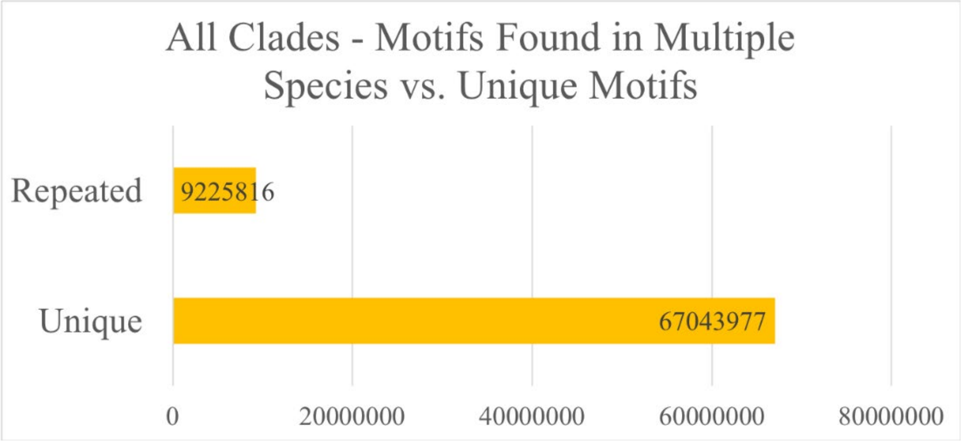


Supplementary Figure 2:

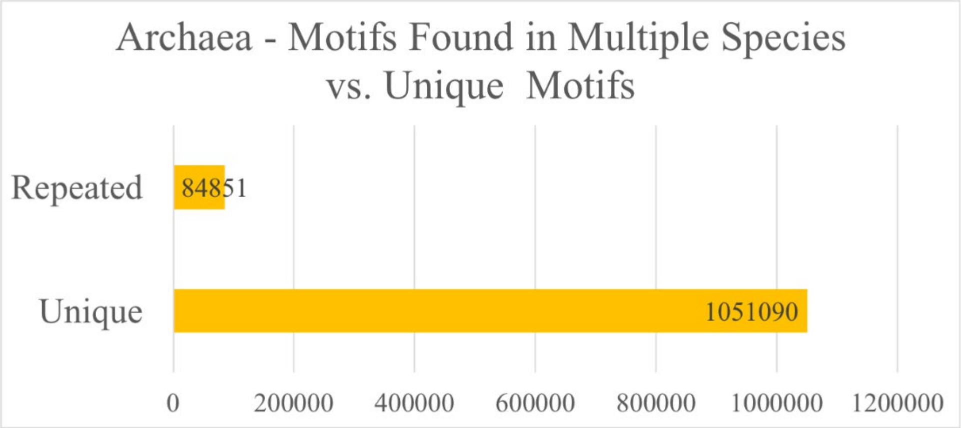

Supplementary Figure 3:

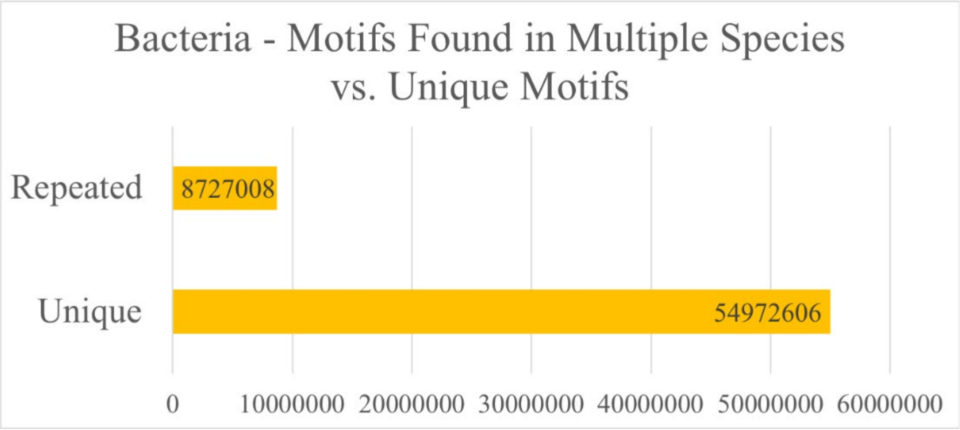


Supplementary Figure 4:

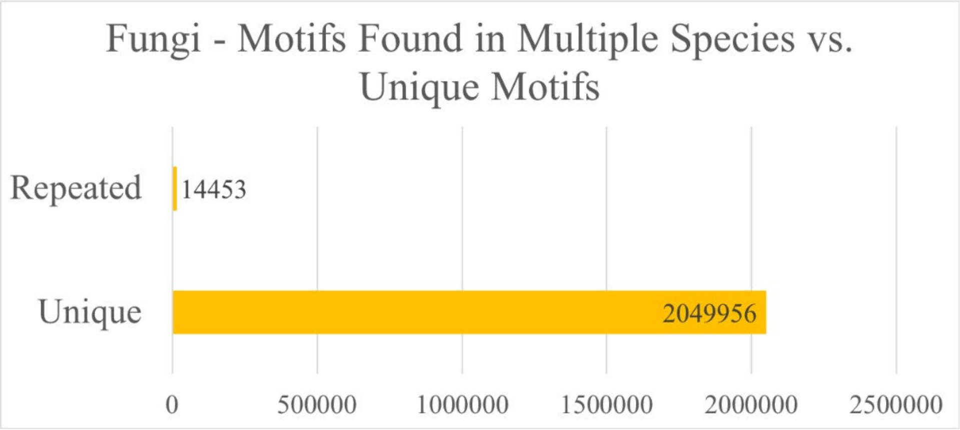


Supplementary Figure 5:

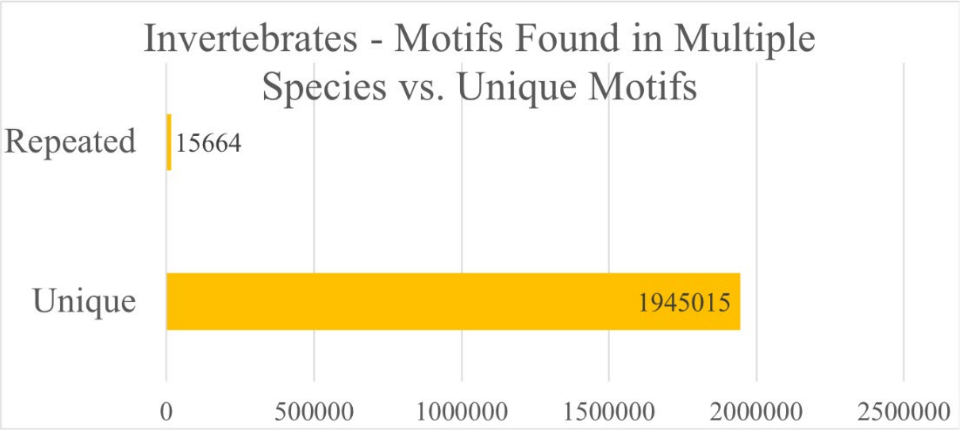

Supplementary Figure 6:

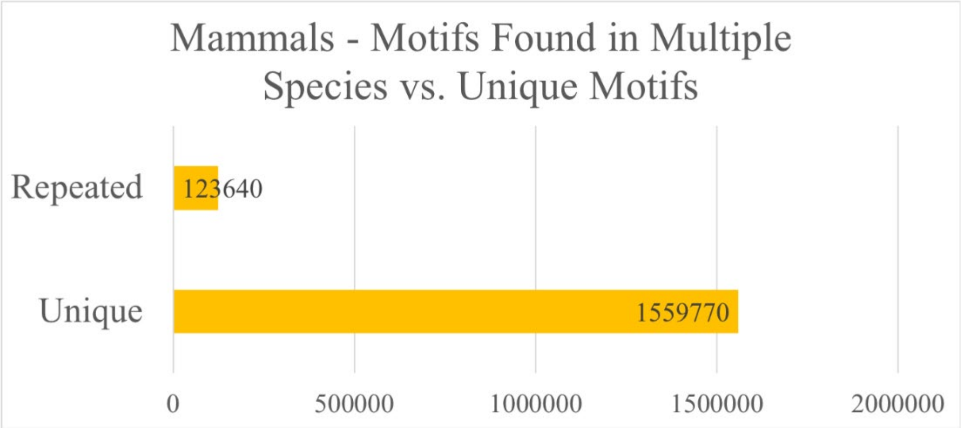


Supplementary Figure 7:

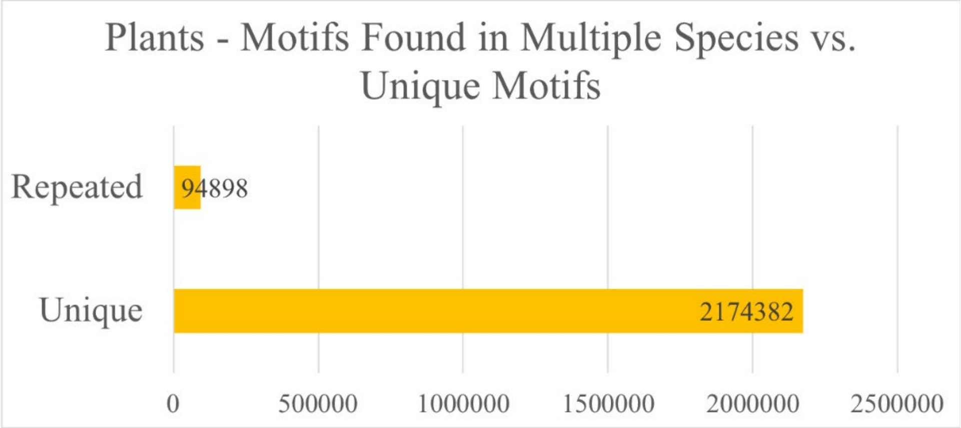


Supplementary Figure 8:

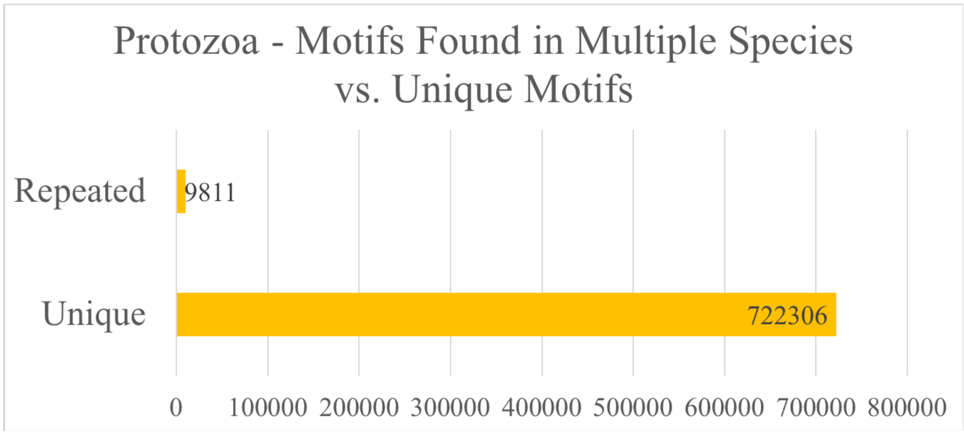

Supplementary Figure 9:

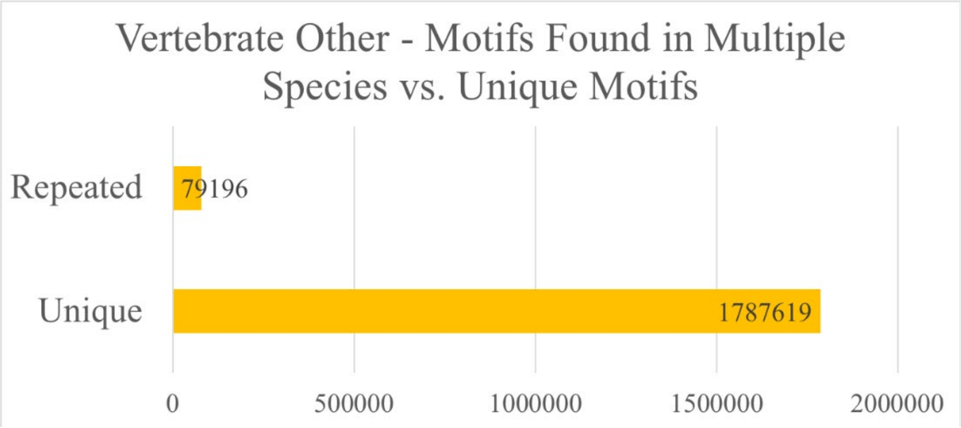


Supplementary Figure 10:

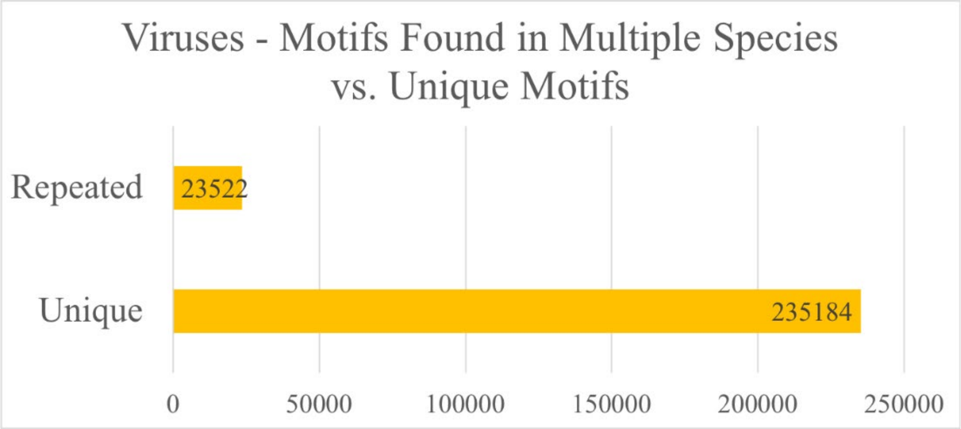


Supplementary Figure 11:

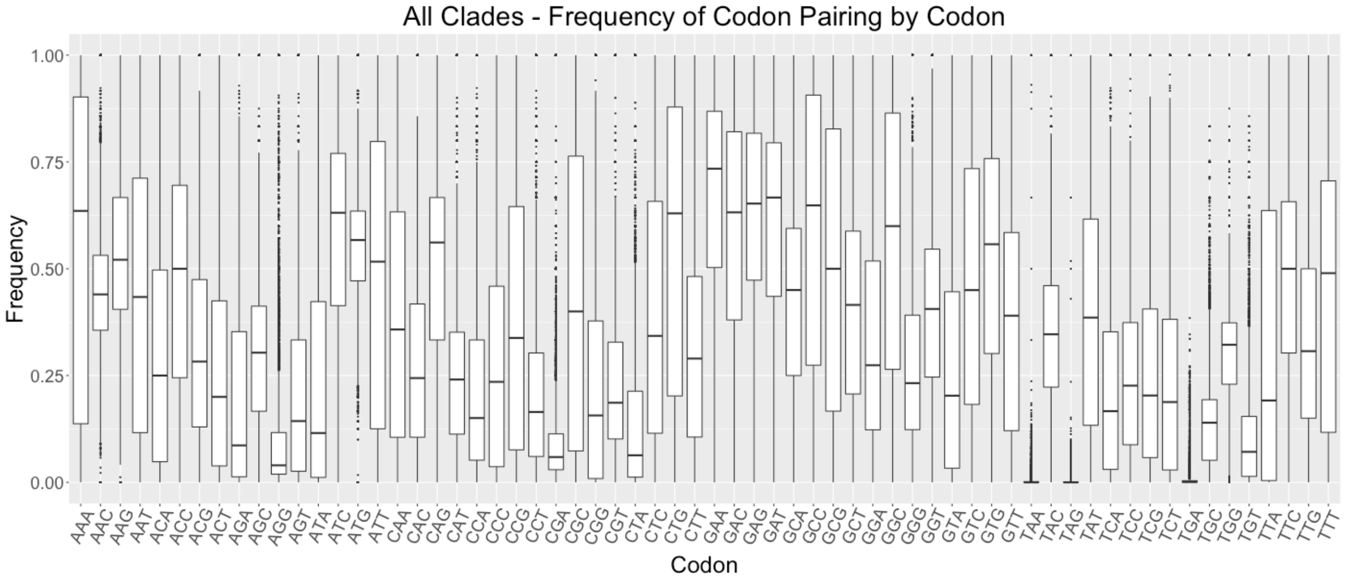

Supplementary Figure 12:

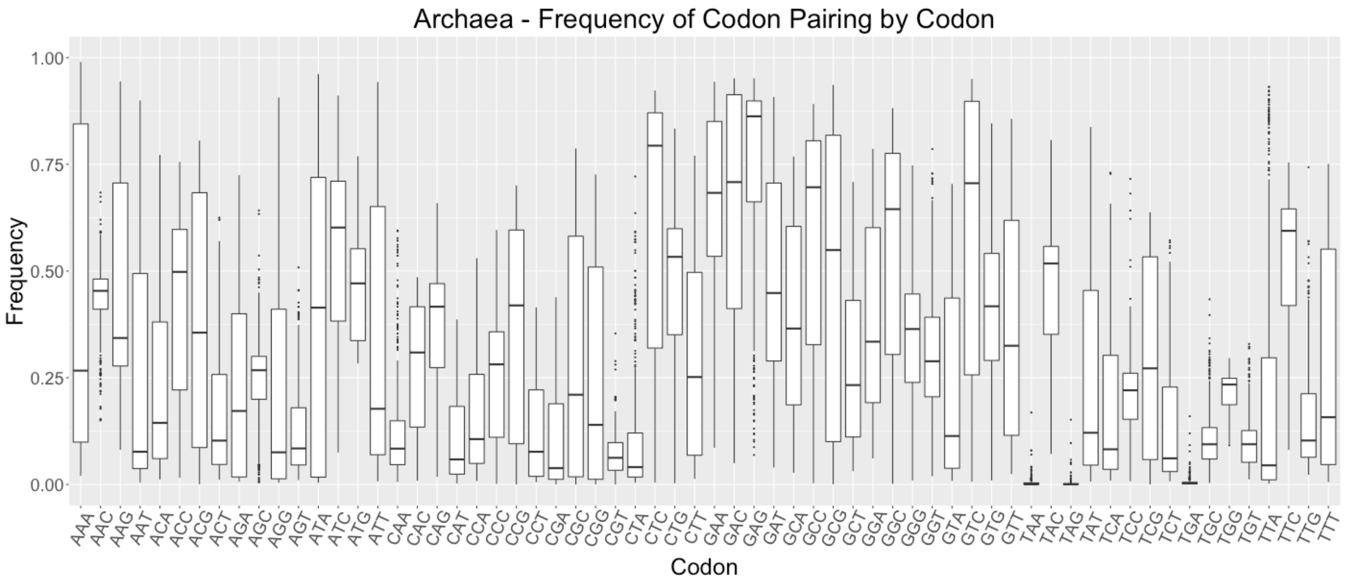


Supplementary Figure 13:

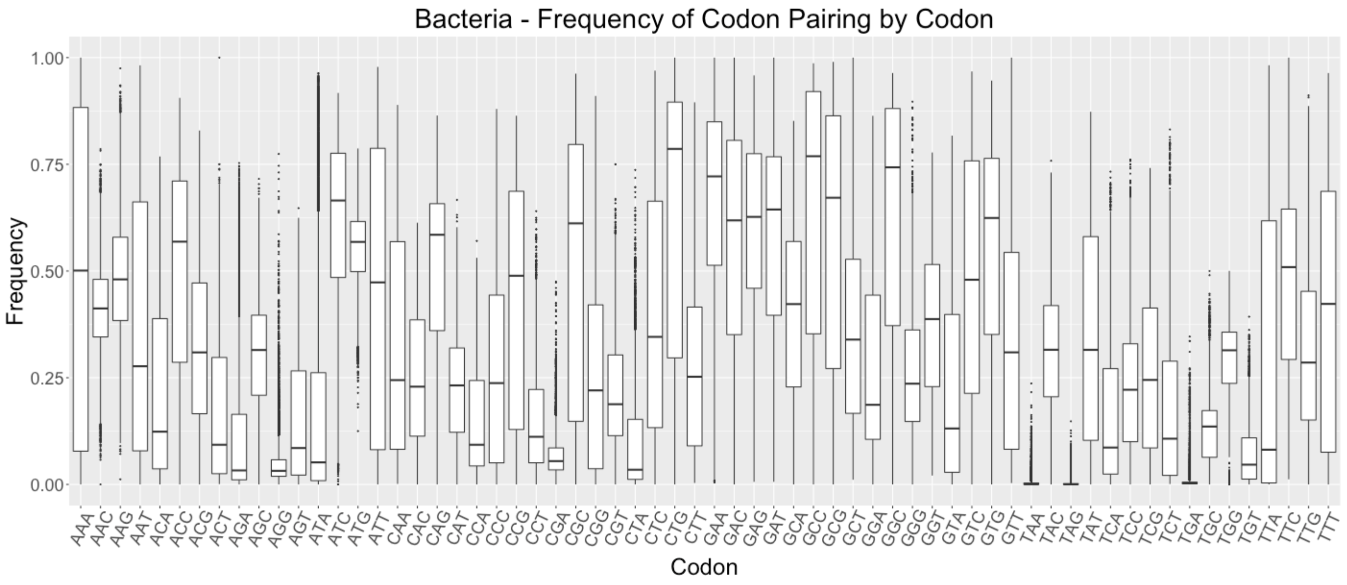

Supplementary Figure 14:

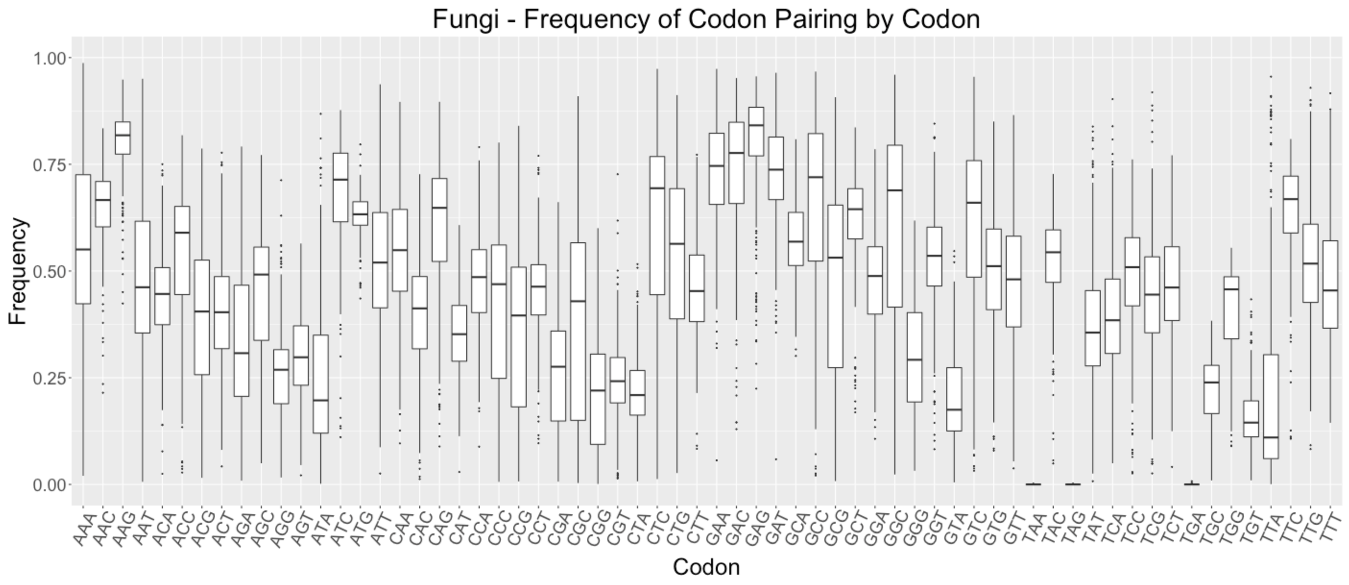


Supplementary Figure 15:

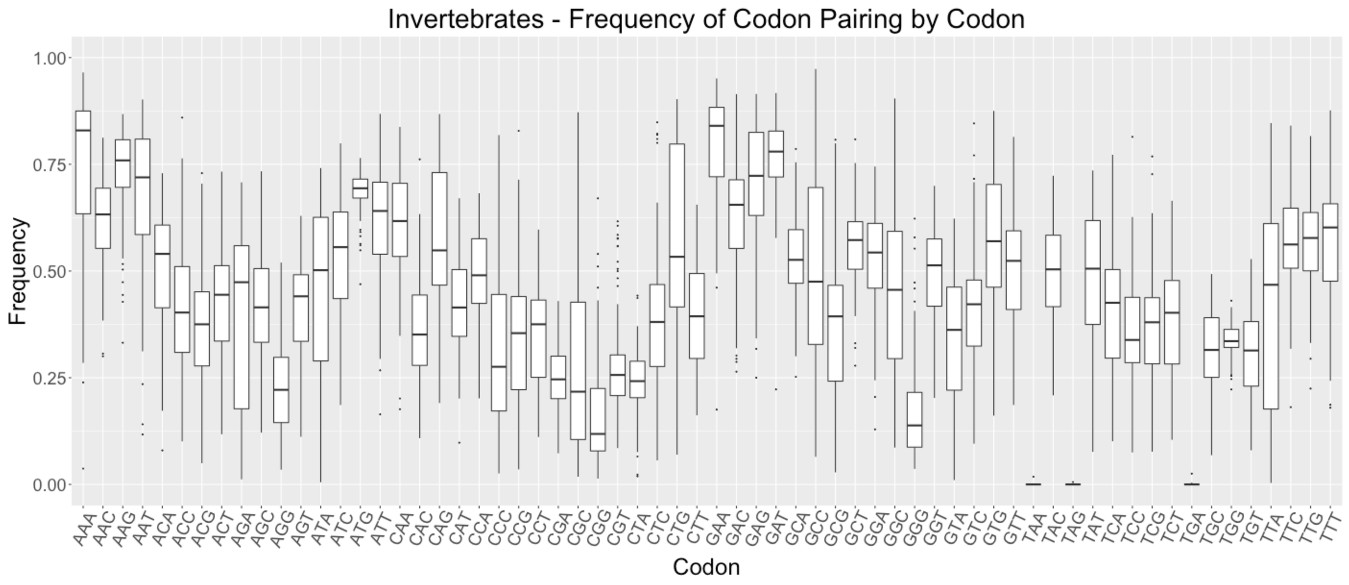

Supplementary Figure 16:

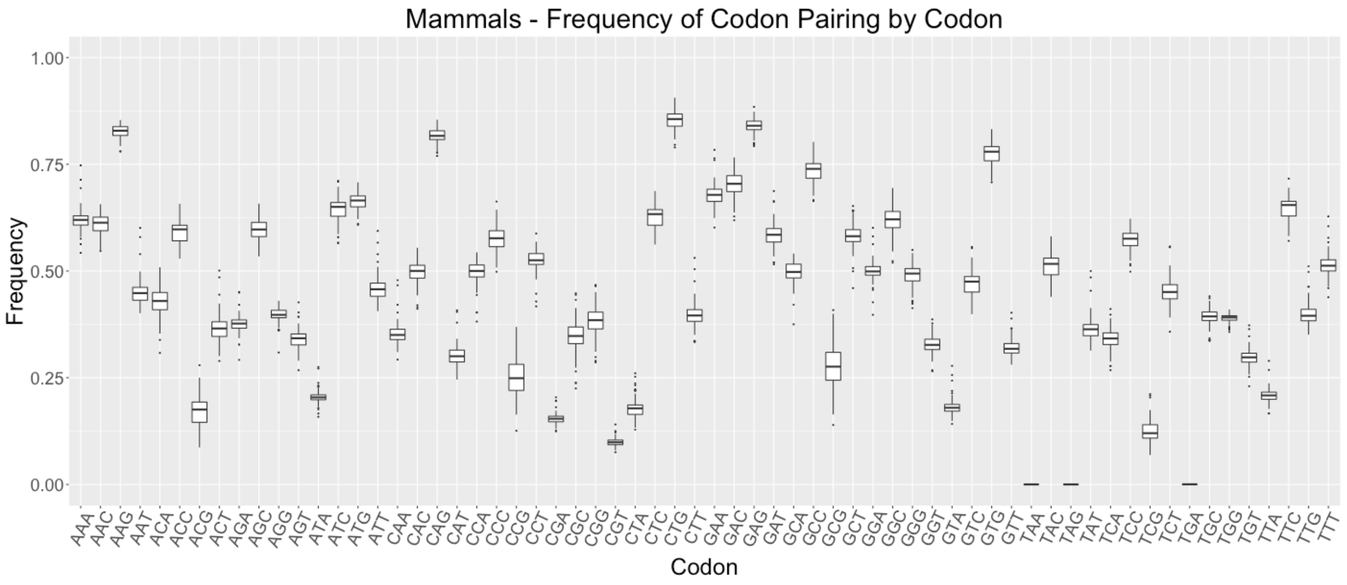


Supplementary Figure 17:

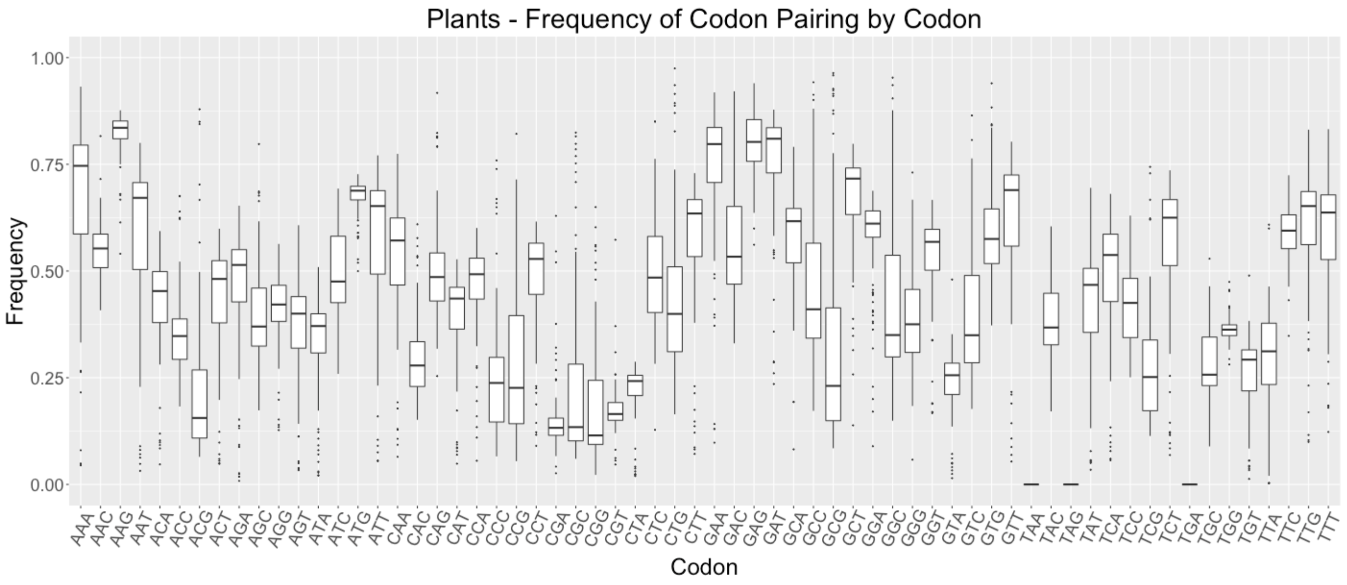

Supplementary Figure 18:

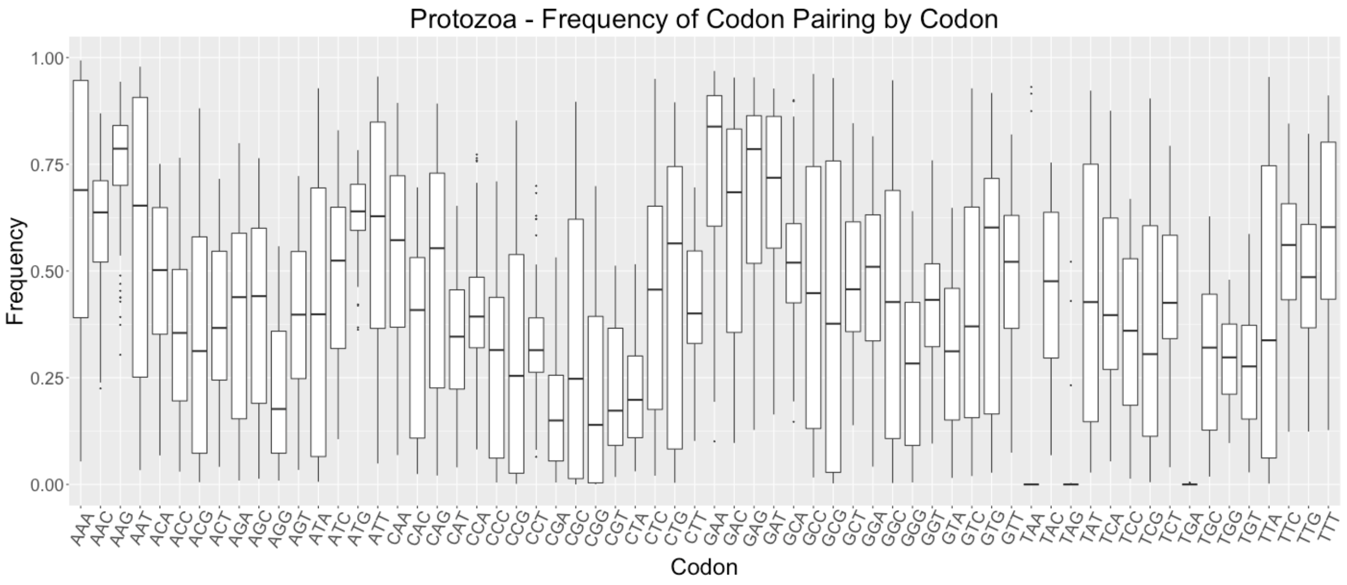


Supplementary Figure 19:

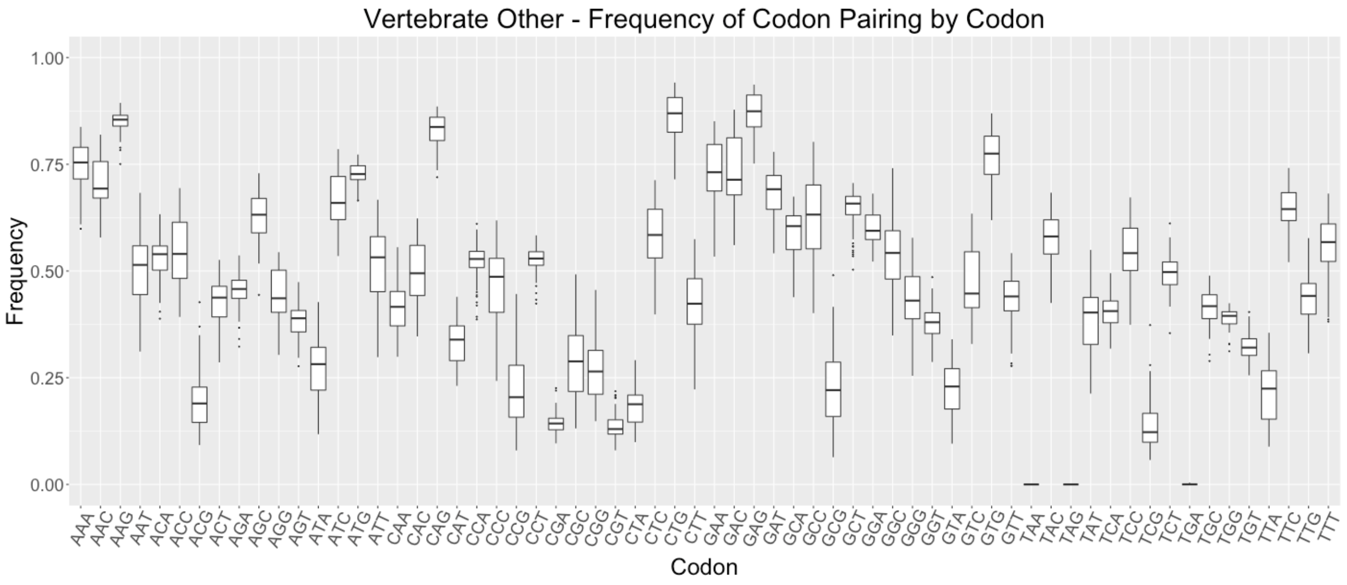

Supplementary Figure 20:

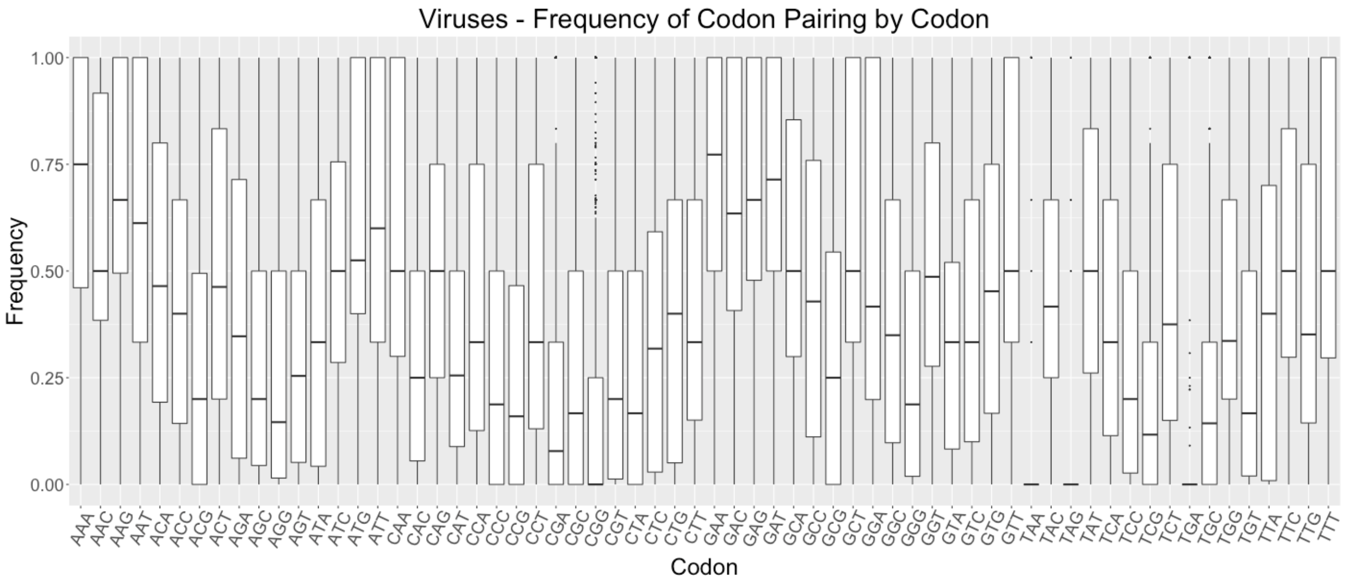


Supplementary Figure 21:

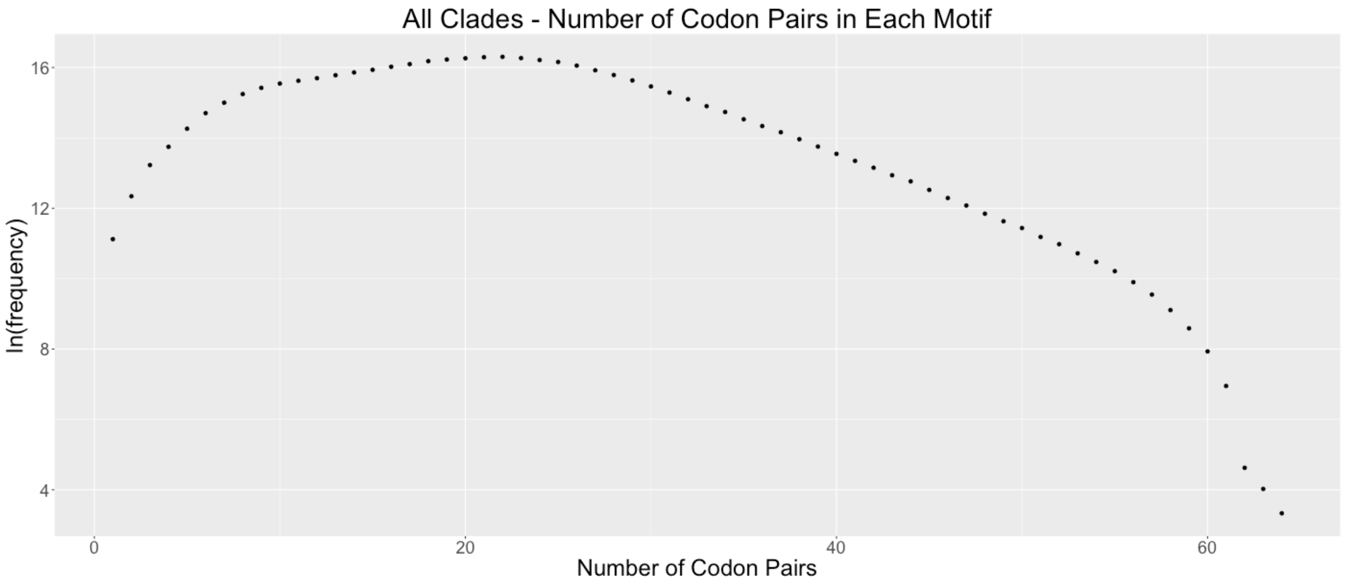

Supplementary Figure 22:

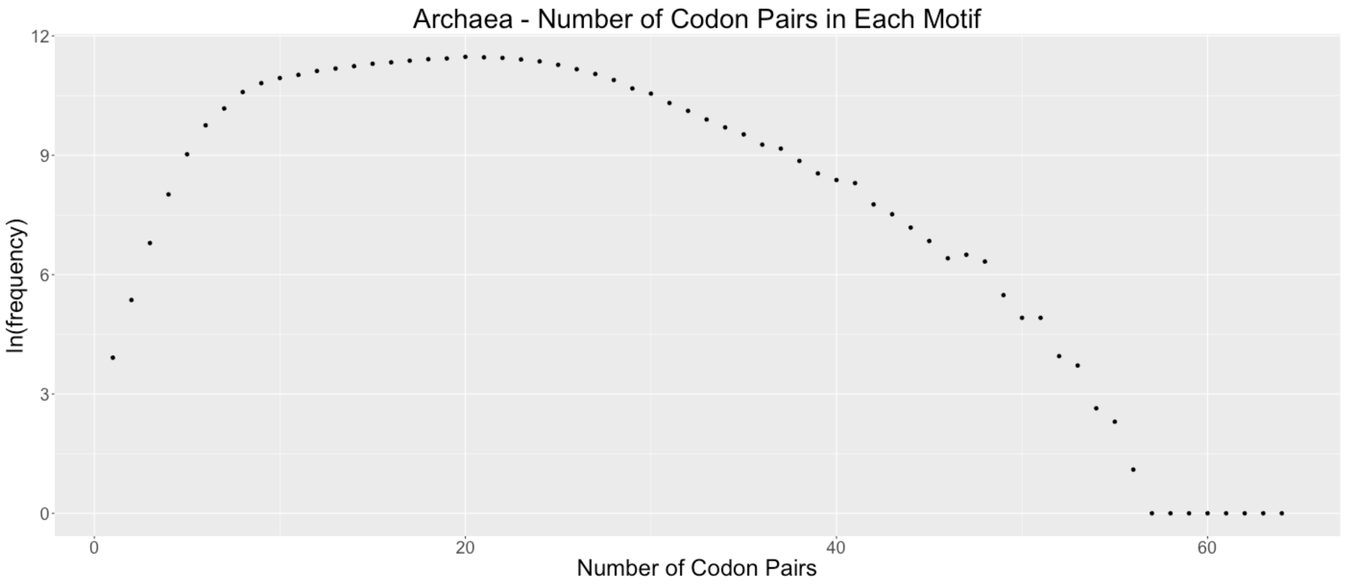


Supplementary Figure 23:

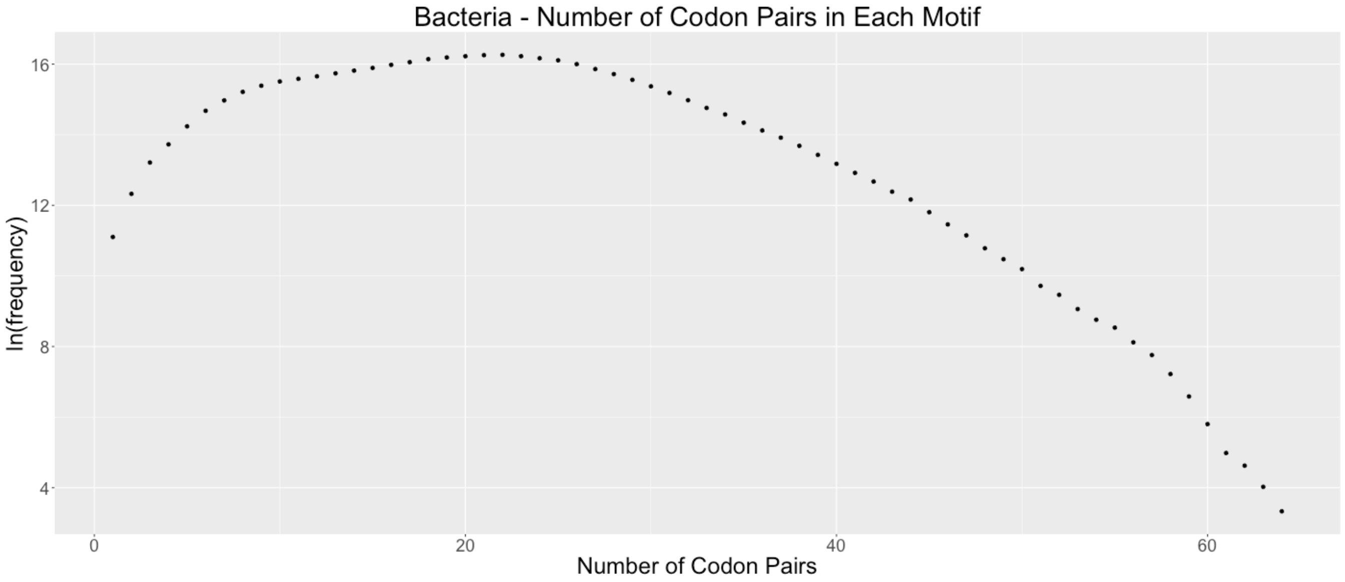

Supplementary Figure 24:

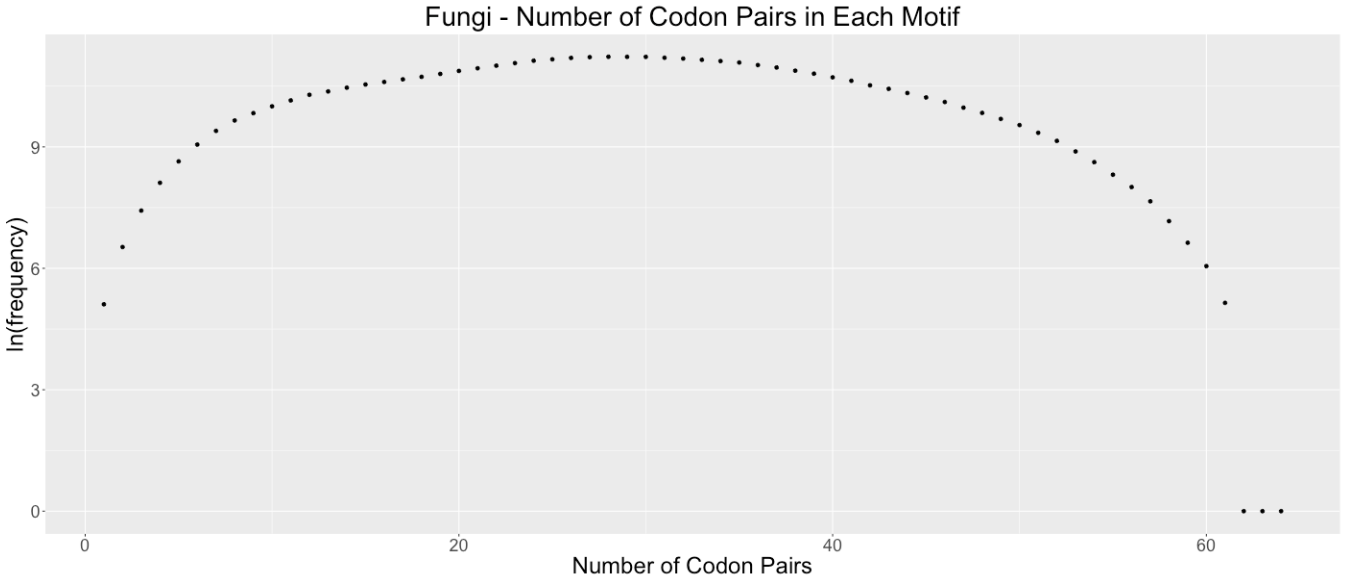


Supplementary Figure 25:

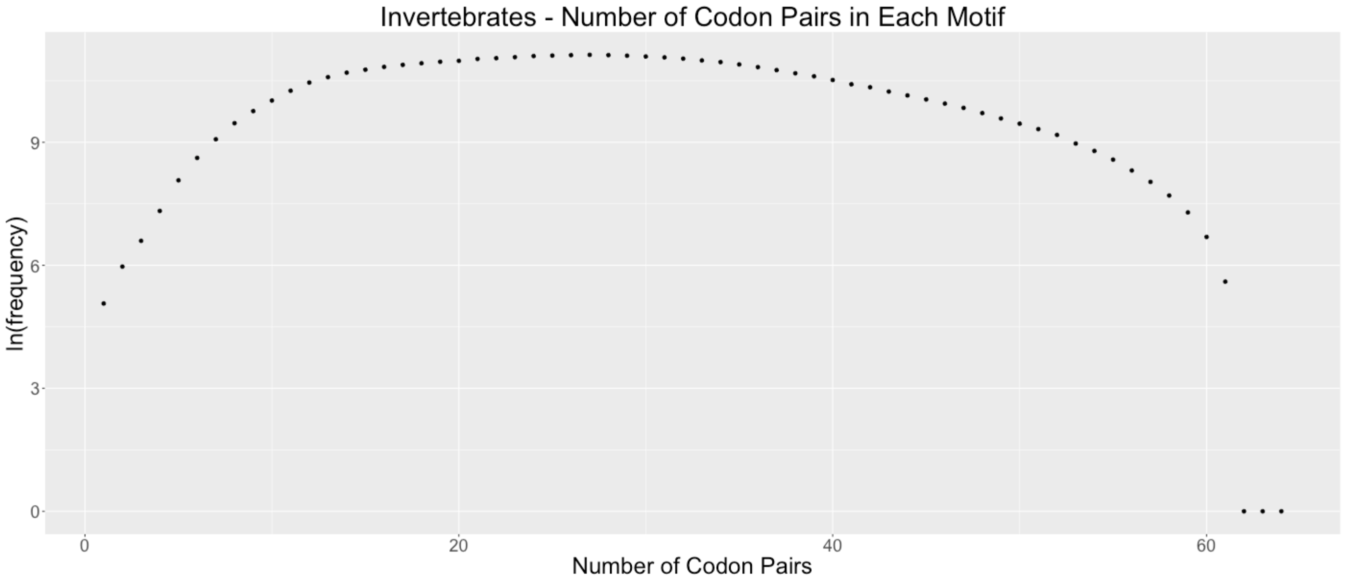

Supplementary Figure 26:

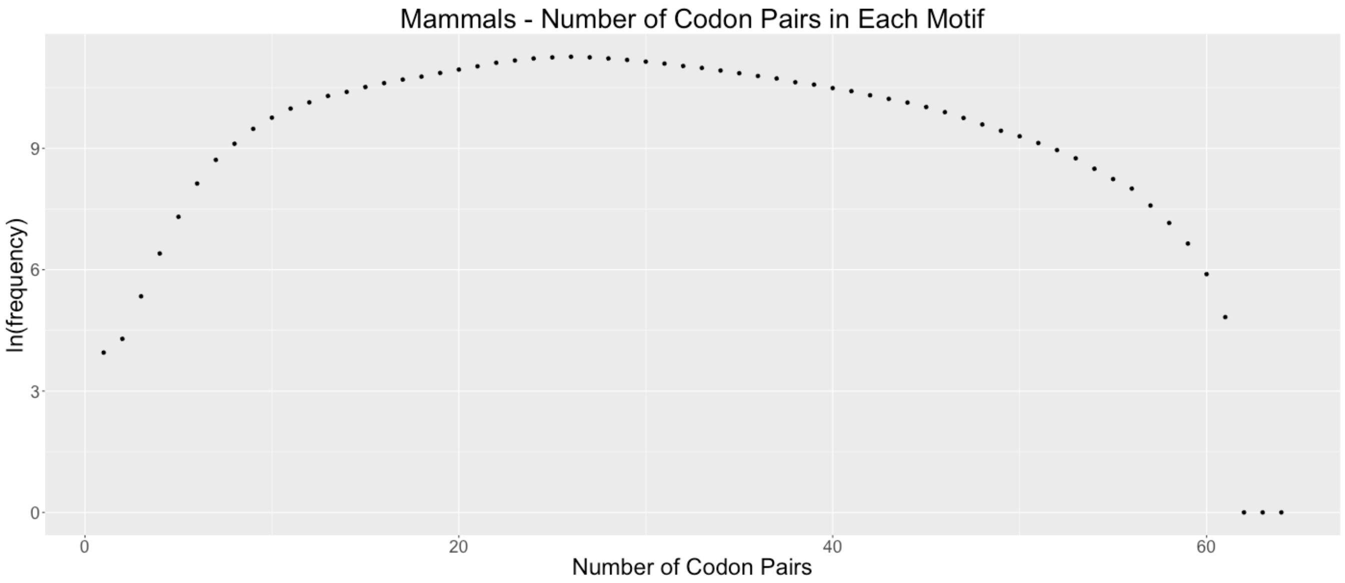


Supplementary Figure 27:

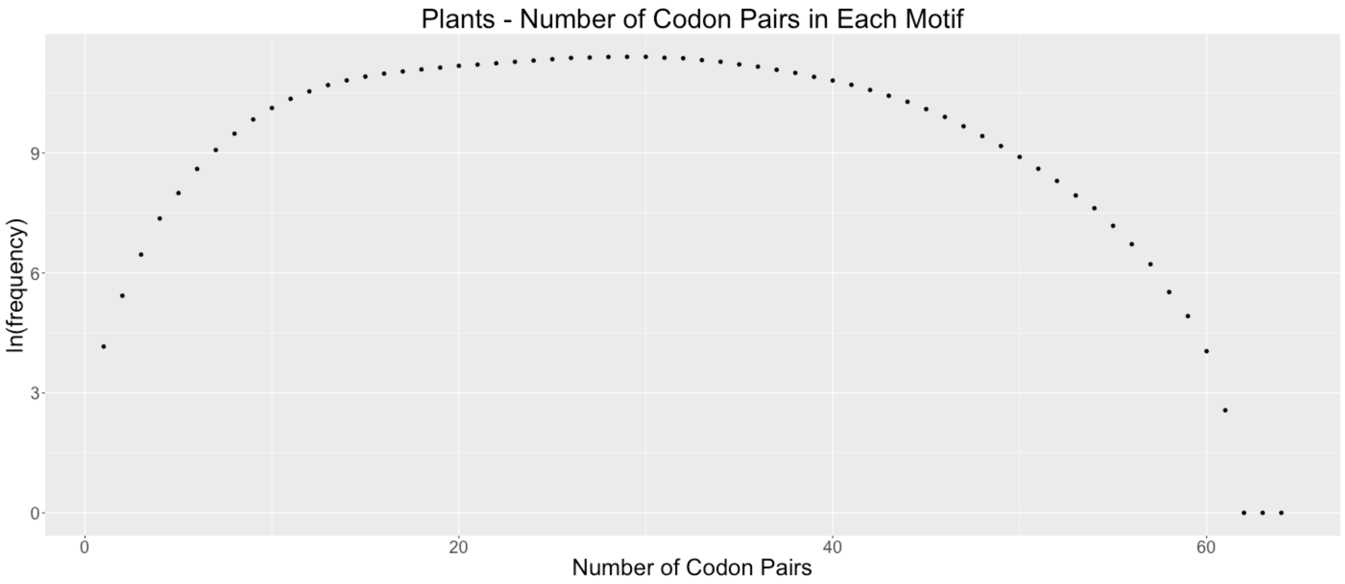

Supplementary Figure 28:

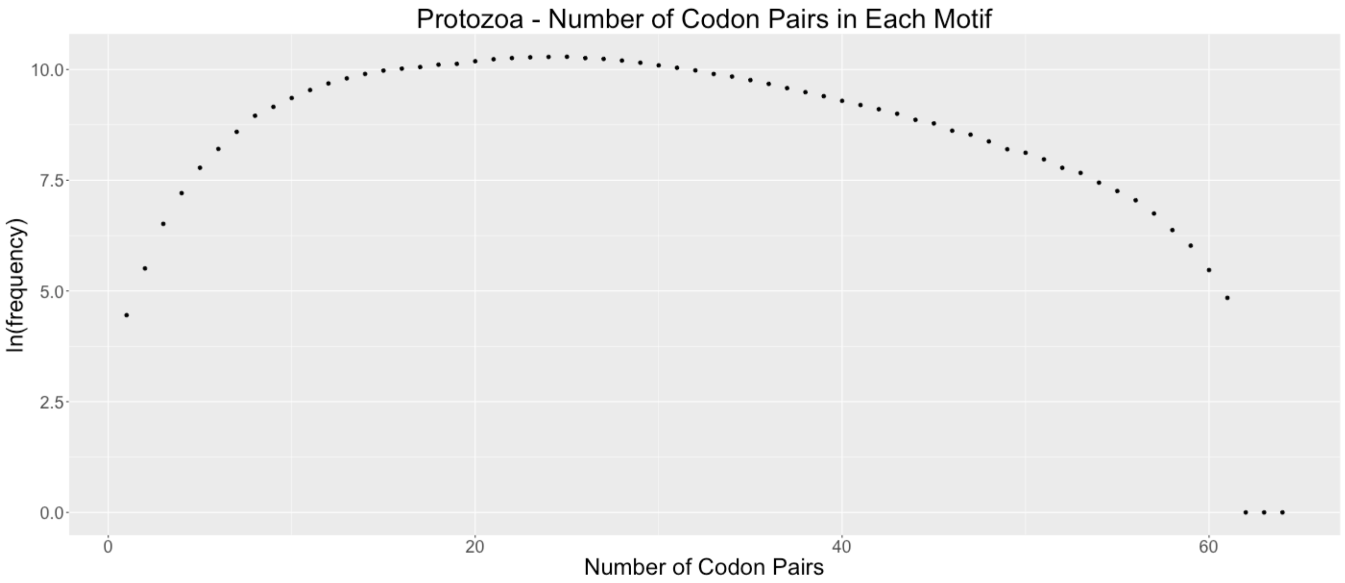


Supplementary Figure 29:

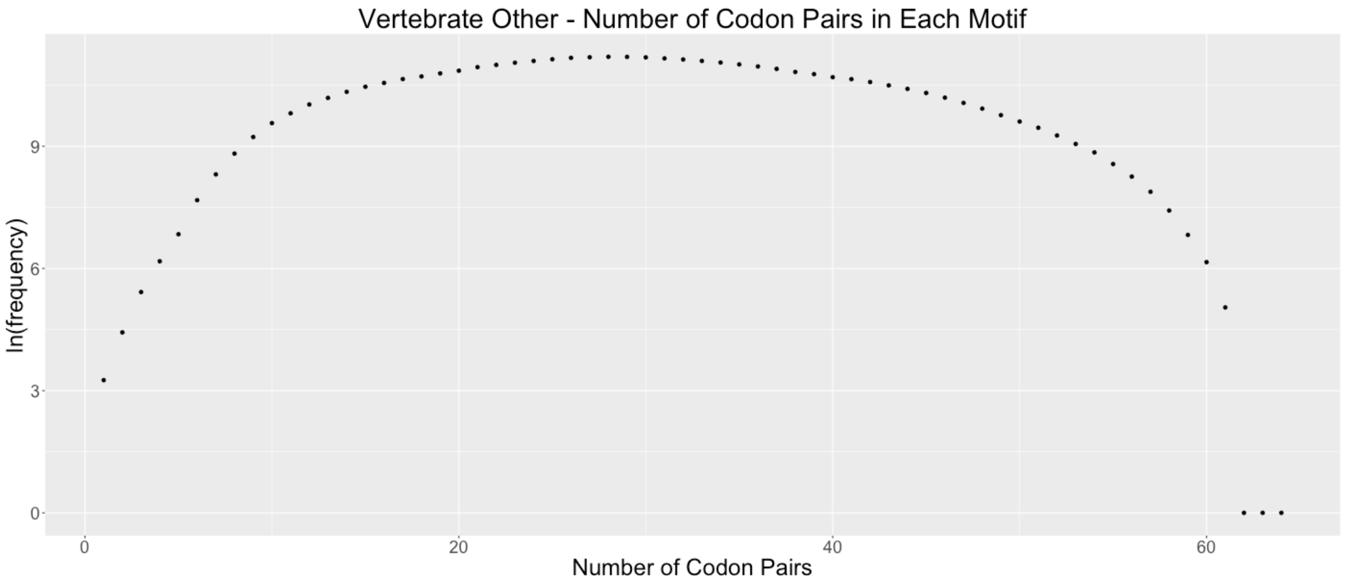

Supplementary Figure 30:

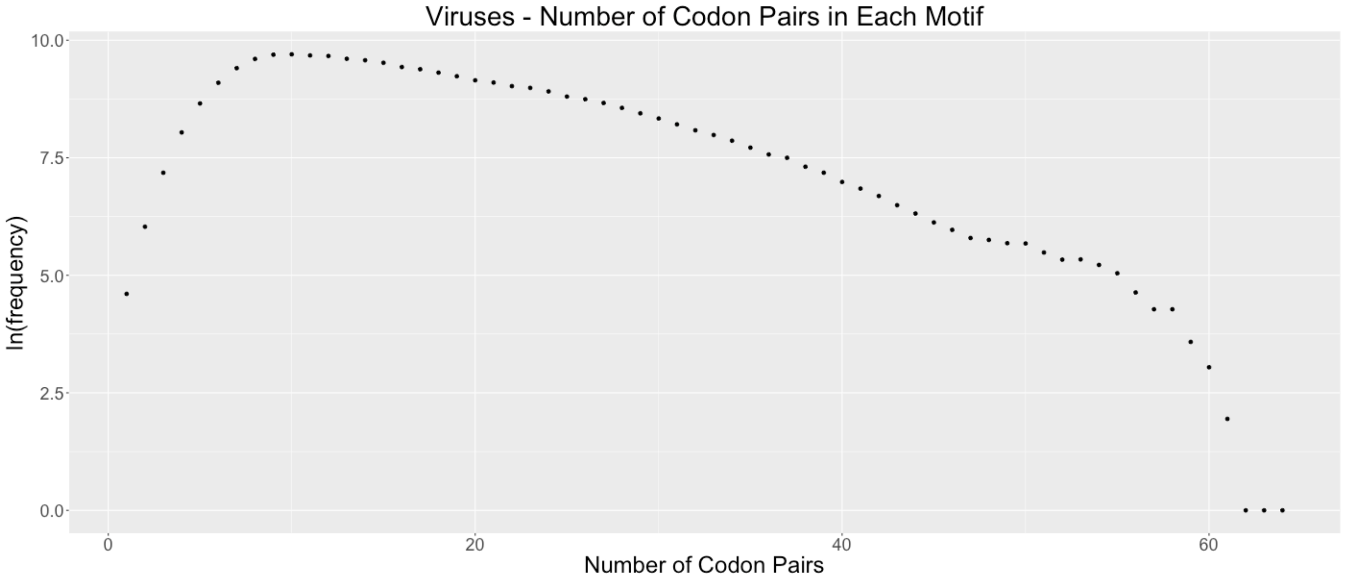


Supplementary Figure 31:

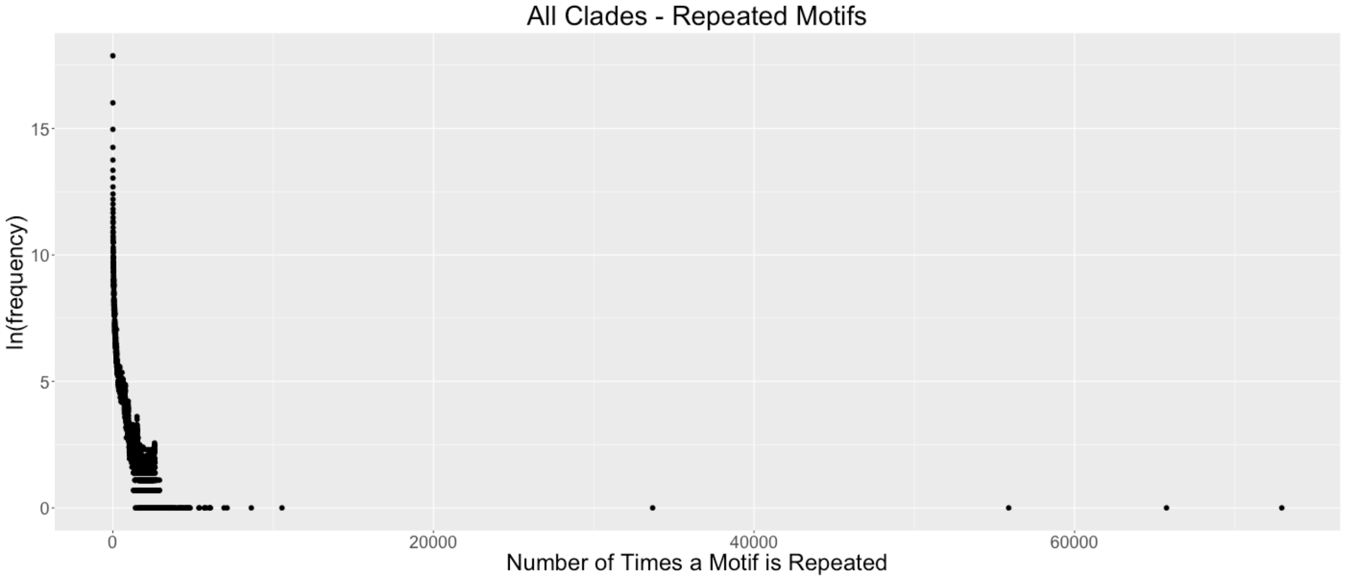

Supplementary Figure 32:

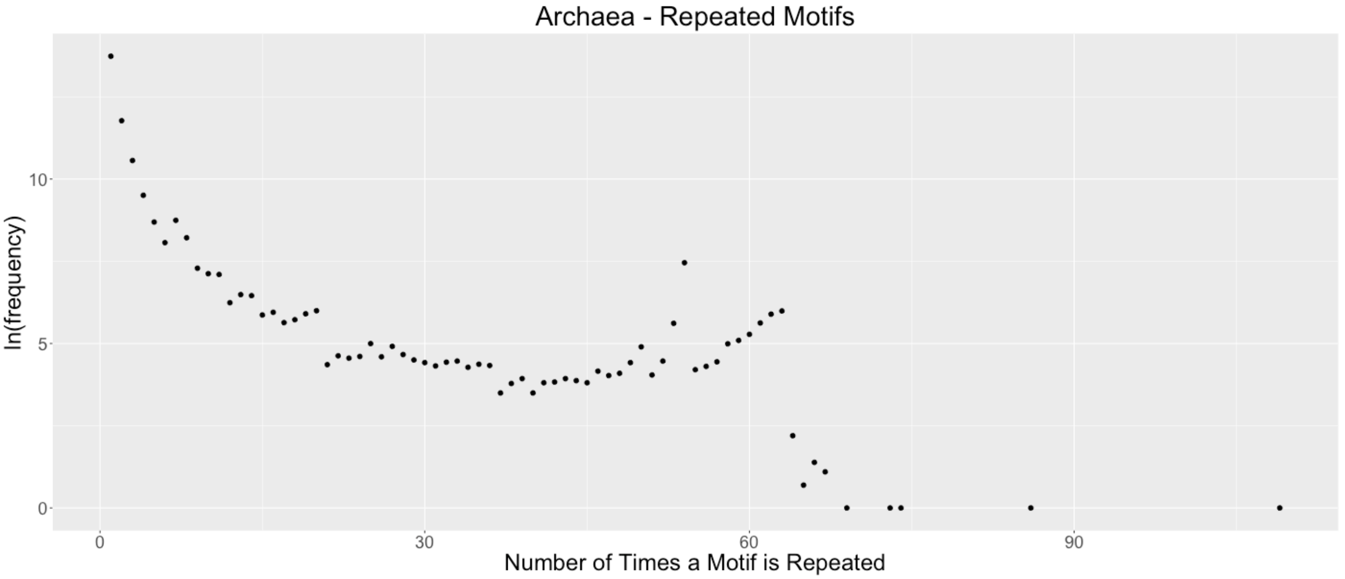


Supplementary Figure 33:

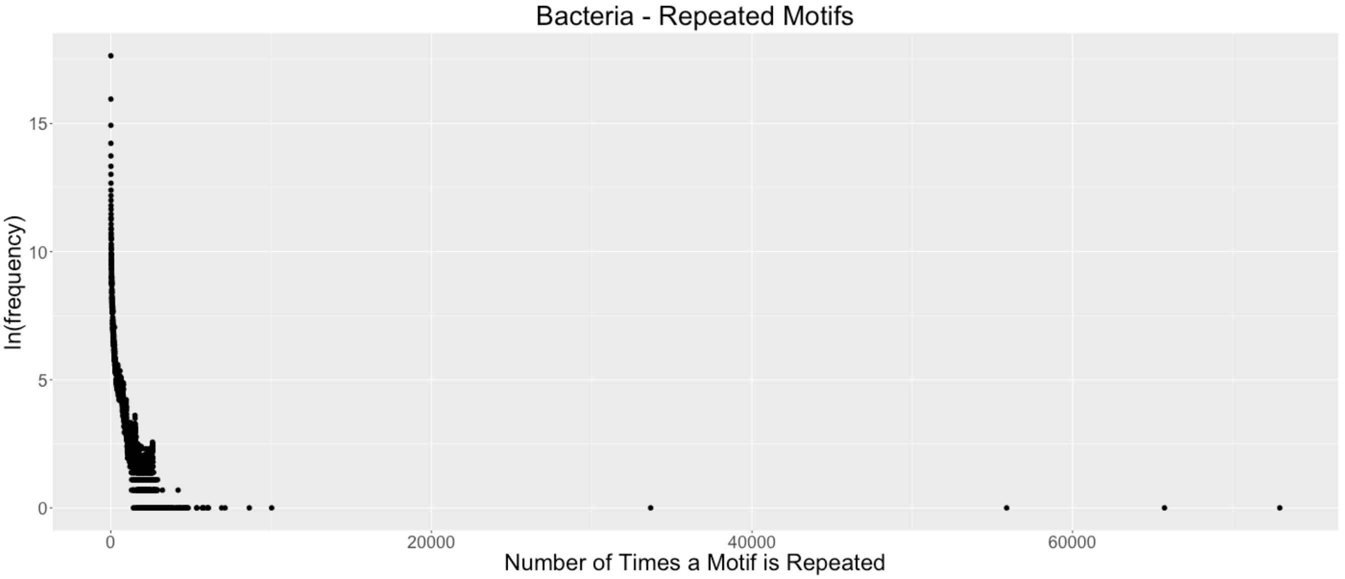

Supplementary Figure 34:

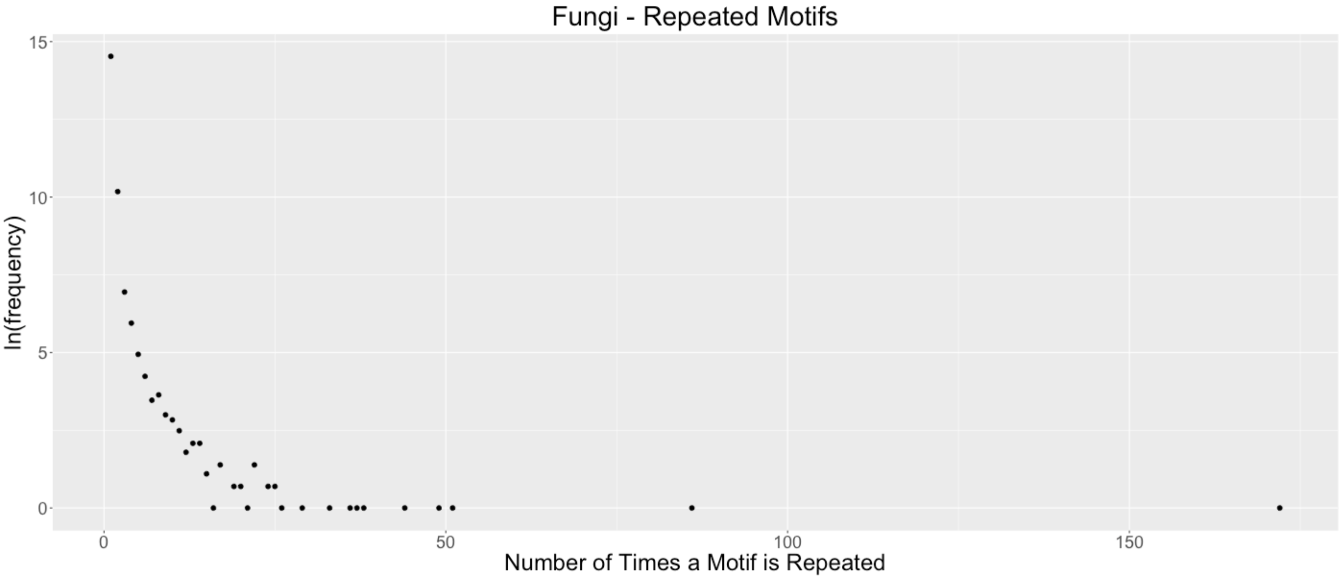


Supplementary Figure 35:

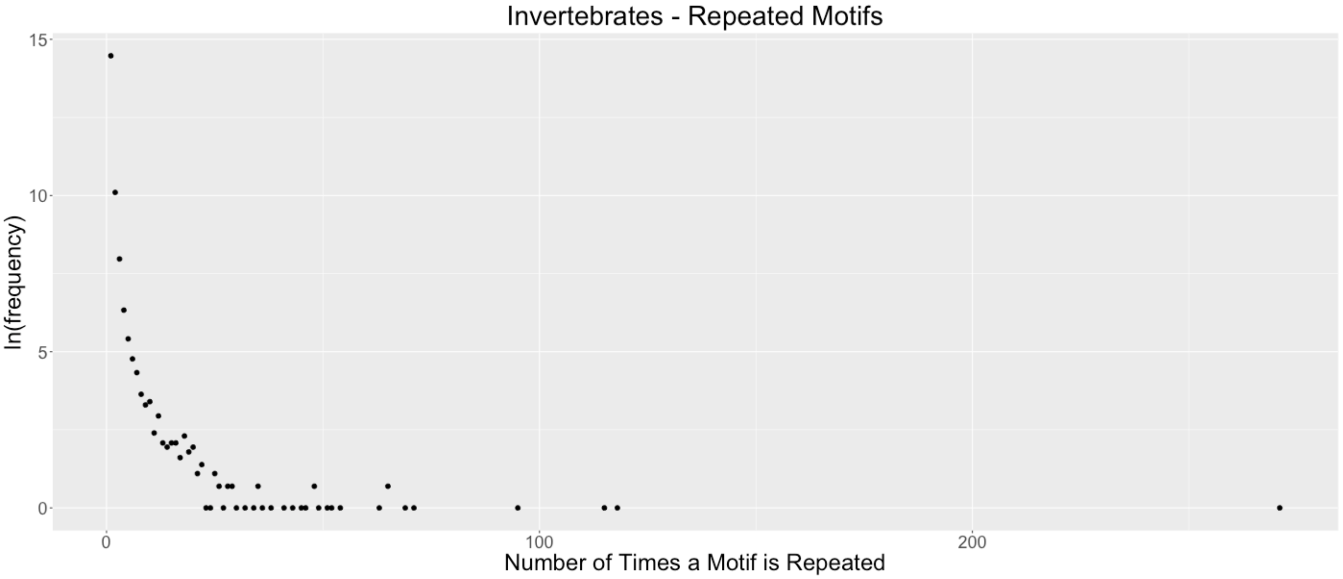

Supplementary Figure 36:

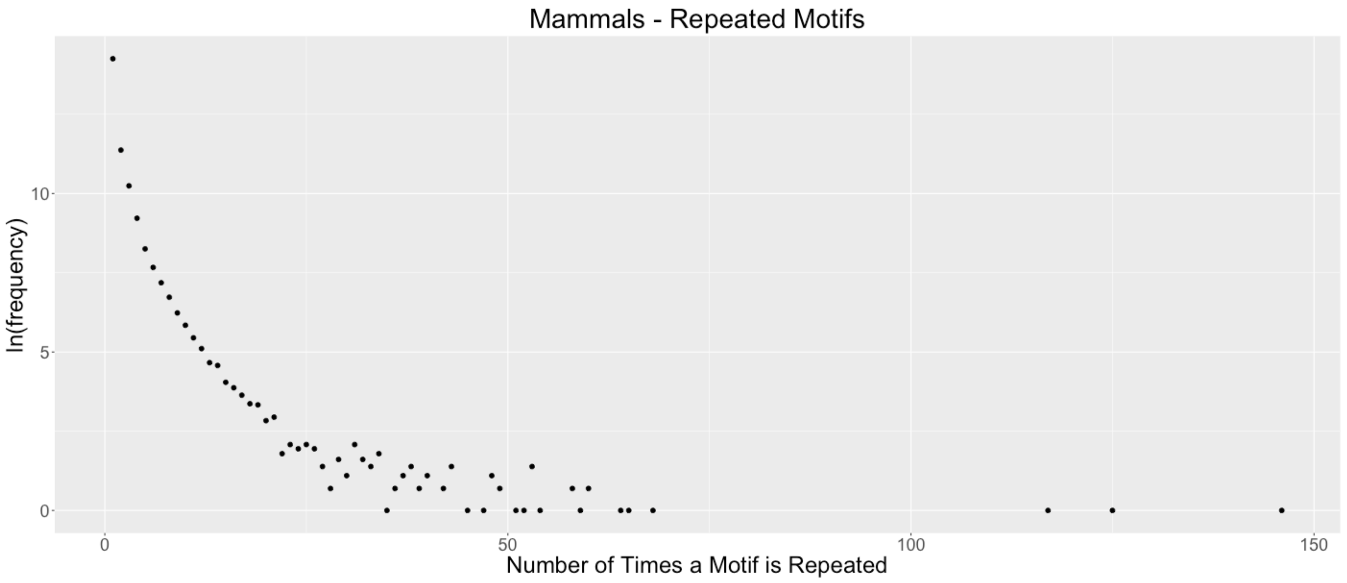


Supplementary Figure 37:

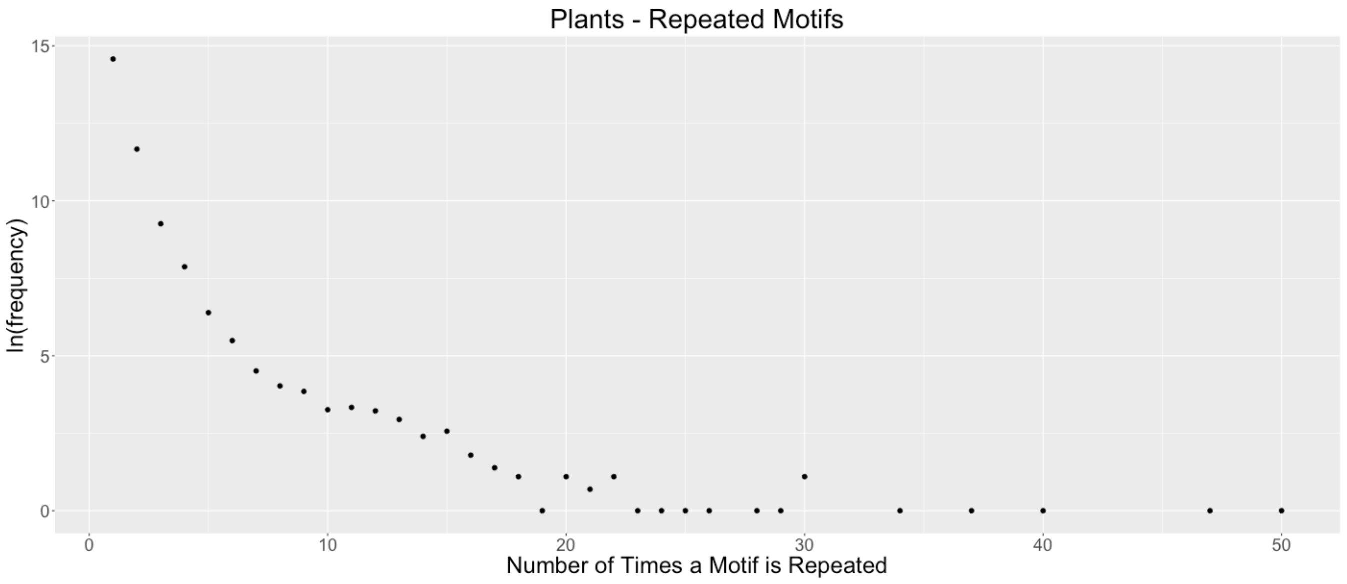

Supplementary Figure 38:

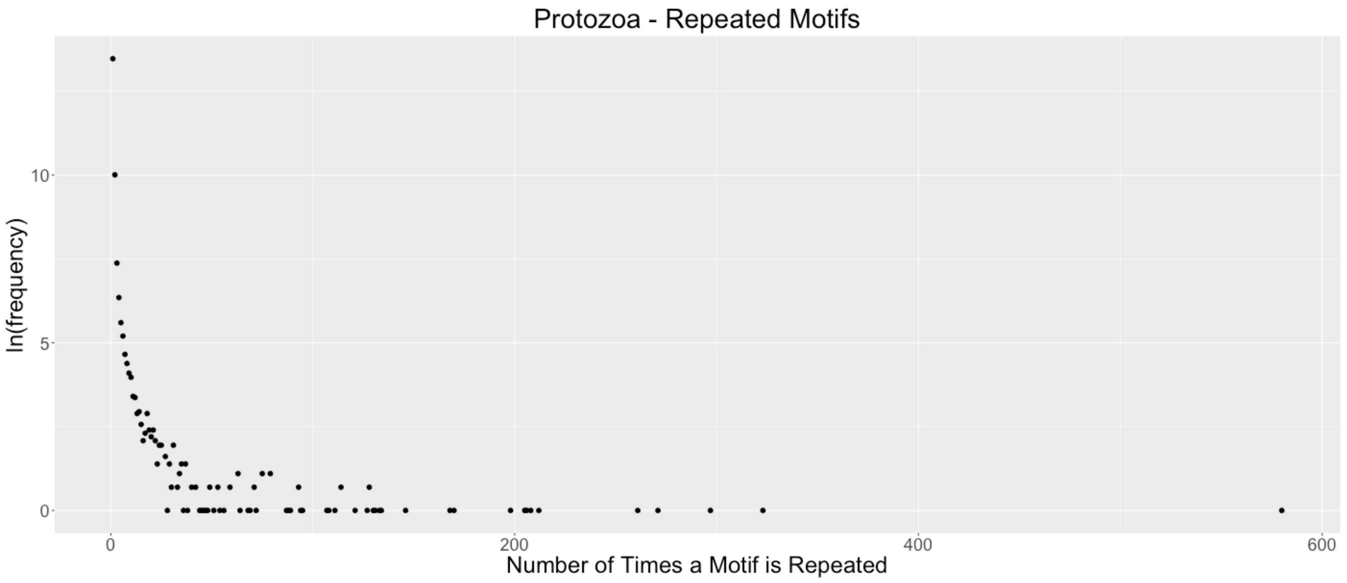


Supplementary Figure 39:

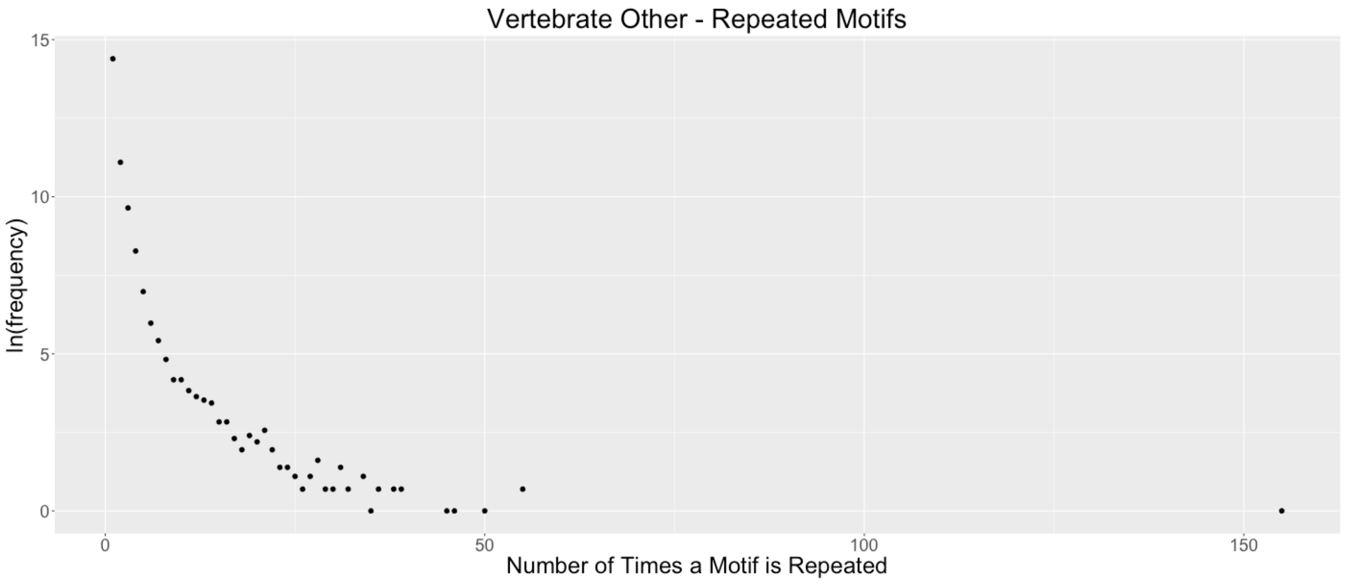

Supplementary Figure 40:

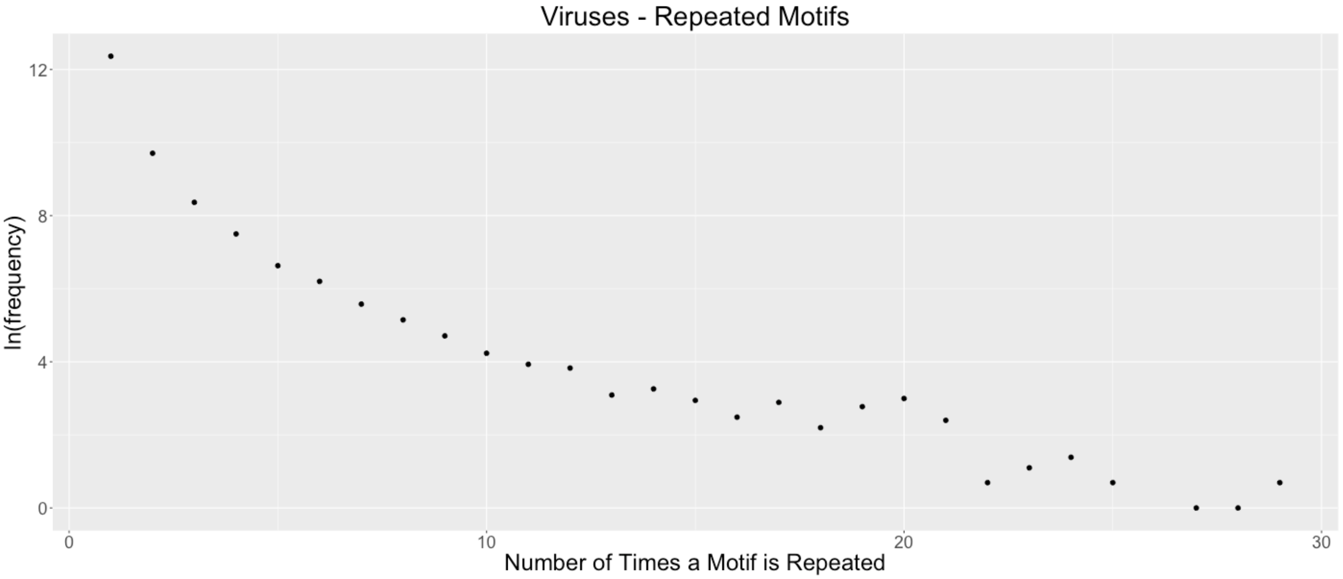
